# Congenital CMV infection drives oligoclonal expansion of cytotoxic γδ T cells from early fetal progenitors

**DOI:** 10.1101/2025.11.20.689355

**Authors:** Justine Levan, Anna V. Vaaben, Jessica Tsui, Brittany R. Davidson, Meagan E. Olive, Perri C. Callaway, Catherine B. Nguyen, Mikias Ilala, Ravi K. Patel, Riana D. Hunter, Gabriela K. Fragiadakis, Alexis J. Combes, Felistas Nankya, Grant Dorsey, Abel Kakuru, Mary Muhindo, Margaret E. Feeney

## Abstract

Gamma delta T cells (γδ T cells) emerge early during human gestation and are uniquely equipped to protect the fetus and infant following *in utero* infection with viruses such as cytomegalovirus (CMV). Previous research showed that fetal γδ T cells in early gestation are transcriptionally pre-programmed for effector functions in the thymus. Infants with congenital CMV infection (cCMV) exhibit expansions of γδ T cells with CMV-reactive TCRs; however, the functional and transcriptional programming of these innate-like effector cells has not been characterized. Here, we analyzed cord blood mononuclear cells from cCMV+ and uninfected neonates in Uganda using flow cytometry and single-cell RNA and TCR sequencing. We find that γδ T cells in cCMV+ neonates are more differentiated, activated, cytotoxic, and proliferative. TCR repertoires of cCMV+ infants exhibit oligoclonal expansions with an enrichment of γδTCRs that possess shorter CDR3 lengths and fewer N additions, which suggest they arise from early fetal progenitor cells. These expanded γδ T cell clonotypes in cCMV+ infants are more frequently public and exhibit cytotoxic transcriptional programming. These findings demonstrate that cCMV infection drives an oligoclonal expansion of highly cytotoxic effector γδ T cells with fetal-like TCR features, underscoring their specialized roles in early-life immunity.

## INTRODUCTION

The human fetus and young infant are uniquely susceptible to infections, as they lack adaptive immune memory from prior encounters with pathogens. Thus, to counter infectious threats, the fetus relies on immune cell populations and differentiation programs that are poised for rapid effector function^1^. T cells expressing the γδTCR are the first to develop during human gestation, appearing in the fetal liver as early as 6-9 weeks^2^. They recognize non-peptide antigens and possess many features that make them uniquely suited to protecting the fetus and infant, including rapid, innate-like effector functions that are not dependent on prior antigen exposure or priming by dendritic cells, which are functionally immature in early life^3–7^. Further, they are MHC-unrestricted and exhibit remarkable functional plasticity, including cytolytic, antibody-mediated, and antigen-presenting functions^8–11^. In some infection models, γδ T cells are required for the protection of young, but not mature, mice^12^. Indeed, it has been hypothesized that the primary selective advantage driving the γδTCR’s over 450,000,000 years of conservation as a third class of V(D)J-recombined antigen receptors is its role in neonatal protection^13^.

Human cytomegalovirus (CMV) is the most common congenital infection worldwide^14^. It is a leading cause of neurodevelopmental delays in newborns, and the clinical consequences of *in utero* infection may be severe, including microcephaly, hepatosplenomegaly, jaundice, chorioretinitis, hearing loss, or being small for gestational age (SGA)^15,16^. In adults, CMV infection induces an expansion of γδ T cells that can lyse CMV-infected target cells *in vitro*^17–22^. The precise ligand(s) targeted by γδ T cells in CMV-infected individuals are largely unknown and may include viral-encoded or stress-induced host cell ligands^23–26^. While specific subsets of γδ T cells have also been observed to expand in congenitally CMV-infected (cCMV) infants, the γδTCR repertoire in such infants is notably restricted, oligoclonal, and highly shared. One study of European infants demonstrated the presence of a highly shared (“public”) Vγ8+Vδ1+ clonotype that was entirely germline-encoded and present in nearly all CMV-infected neonates^27^. In contrast, CMV+ adults possess diverse and largely private repertoires^22,28^. The observed differences between the adult and fetal γδTCR cell repertoires upon infection with CMV may be explained by the “layered” model of immune development, which posits that the immune system develops via successive waves of T cells that arise from distinct progenitor populations differing in their intrinsic properties and developmental potential^1,29–31^. During murine development, γδ T cells emerge in distinct waves that are distinguishable by their Vγ and Vδ chain usage; these subsets arise sequentially and populate the periphery to assume functionally specialized roles^32,33^. Recent data indicate that human γδ T cells also emerge in discrete waves that derive from distinct hematopoietic precursor populations and exhibit differences in antigen recognition, functional programming, and other intrinsic properties^31,34–37^. In humans, however, these developmental waves are not as easily defined by Vγ or Vδ chain usage, yet detailed analyses of γδTCRs of mid-gestation human thymocytes have revealed distinct sequence characteristics associated with early fetal origin, including limited junctional diversity and preferential gene segment usage^34^. Importantly, these mid-gestation γδ T cells exhibit effector-biased transcriptional programming, even prior to thymic egress or encounter with pathogens^34^.

Given their unique significance in early life immunity, we sought to determine how γδ T cells in the developing fetus respond to congenital CMV infection. Here, we employ flow cytometry and paired single-cell transcriptional and TCRseq profiling of individual cord blood γδ T cells from cCMV-infected and uninfected Ugandan infants to examine how *in utero* infection with CMV influences the phenotype, gene expression, TCR repertoire, and effector functions of γδ T cells in the fetus. Our findings suggest that congenital CMV infection triggers an oligoclonal expansion of γδ T cells characterized by high expression of cytotoxic granule-associated molecules and the persistence of a distinct fetal transcriptional signature.

## RESULTS

### CMV infection *in utero* results in expansion, effector differentiation, and proliferation of multiple γδ T cell subsets

To characterize the γδ T cell response to congenital CMV (cCMV) infection, we performed flow cytometric analysis on CBMCs collected from 12 cCMV+ and 73 cCMV-neonates from Eastern Uganda. We labeled γδ T cells using antibodies recognizing the Vδ1, Vδ2, and Vγ9 TCR chains and evaluated the frequency of four major subsets: Vγ9-Vδ1+, Vγ9+Vδ1+, Vγ9-Vδ2+, and Vγ9+Vδ2. Among both cCMV-infected and cCMV-uninfected neonates, Vγ9-Vδ1+ cells represented the largest population of γδ T cells analyzed, comprising from 0.34% to over 3% of all T cells **(Figure 1A)**. Neonates with cCMV infection had significantly higher frequencies of Vγ9-Vδ1+, Vγ9+Vδ1+, and Vγ9-Vδ2+ γδ T cells compared to uninfected neonates, but similar frequencies of Vγ9+Vδ2+ cells **(Figure 1A)**. These data align with previous reports that both Vδ1+ and Vγ9-Vδ2+ T cells expand in response to CMV infection in adult solid organ transplant recipients^18,19^ and align with more limited data from congenitally infected infants^27^. Interestingly, staining with Ki67 indicated that proliferation of all four T cell subsets was higher in cCMV+ infants, including the Vγ9+Vδ2+ cells, which were not increased in frequency **(Figure 1B)**. This suggests that while only particular γδ T subsets expand in response to cCMV infection, the infection may create an immune milieu that promotes *in vivo* proliferation of all γδ T cells.

**Figure 1.**
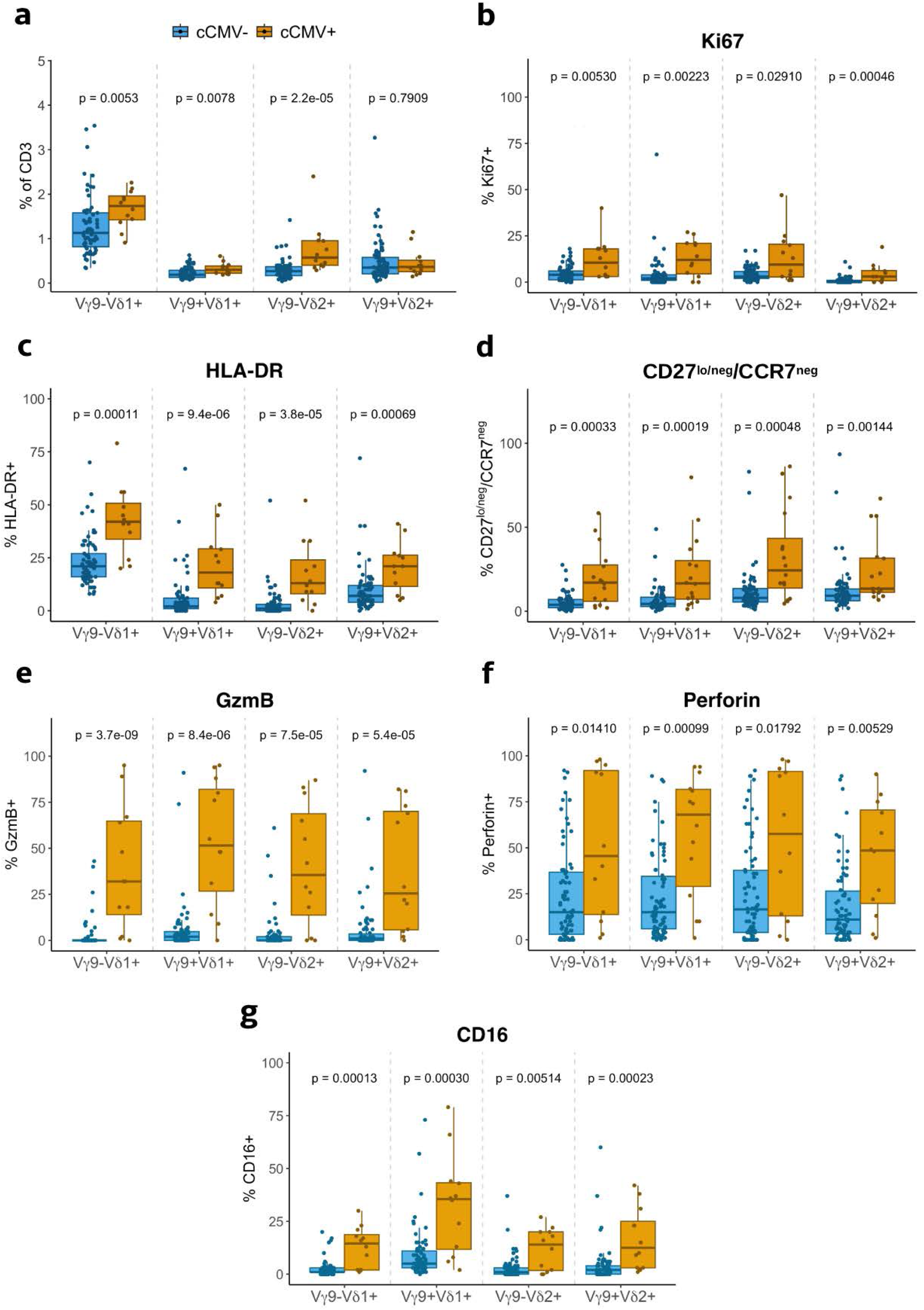
γδ T cells expand and differentiate in response to cCMV. **A)** Quantification of γδ T cell subsets in cord blood of cCMV- (blue; n = 73) or cCMV+ (orange; n =12). Frequency of Ki67+ (**B**), HLA-DR+ (**C**), CD27^lo/neg^/CCR7^neg^ **(D)**, granzyme B **(E)**, perforin **(F)**, or CD16+ **(G)** cells in Vγ9-Vδ1+, Vγ9+Vδ1+, Vγ9-Vδ2+, or Vγ9+Vδ2+ γδ T cell subsets in cCMV- or cCMV+ infants. All flow plots were gated on live CD14^−^CD19^−^CD3^+^ T cells. P-values were determined by Wilcoxon rank sum test.

We next investigated whether these higher frequencies of γδ T cell subsets were accompanied by changes in activation and differentiation. We found that cCMV-infected neonates had a higher frequency of activated, HLA-DR+ cells in all γδ T subsets analyzed **(Figure 1C)**. We then assessed memory differentiation by staining for CD27 and CCR7. Previous publications have defined γδ T cell differentiation based on CD27 expression, with high CD27 expression being associated with naïve cells and low or negative CD27 expression being associated with differentiated cells that are more responsive to short-term stimulation^38^. CD27/CCR7 co-expression is observed on naïve cells present at high frequencies in early life, whereas CCR7^neg^ γδ T cells produce more cytokines^39,40^. Using this framework **(Supplementary Figure 1B)**, we found that across all four γδ T subsets, cCMV+ neonates had significantly greater proportions of CD27^lo/neg^/CCR7^neg^ cells, suggesting greater differentiation. Additionally, within these differentiated CD27^lo/neg^/CCR7^neg^ cells, cCMV-infected neonates exhibited a more dramatic loss of CCR7, indicating a shift in the trafficking of these cells towards peripheral tissues rather than recirculation back to the lymph node.

To further understand the functional differentiation of γδ T cells in response to *in utero* CMV infection, we measured expression of cytotoxic molecules granzyme B (GzmB) and perforin, as well as the low-affinity IgG-Fc receptor CD16. We observed dramatically upregulated expression of GzmB and perforin in cCMV+ neonates across all γδ T subsets **(Figures 1E and 1F)**. This upregulation was especially notable for GzmB, which was rarely expressed by γδ T cells from uninfected neonates, while in cCMV+ neonates, expression was seen in up to 95% of cells. In addition, cCMV-infected neonates displayed significant upregulation of CD16 **(Figure 1G)**, which could enable TCR-independent killing of antibody-opsonized CMV-infected cells^9^.

We also explored whether clinical and demographic parameters influenced the frequency or phenotype of the observed γδ T cell subset expansions **(Supplementary Table 4)**. The frequencies of Vδ1+ and Vγ9-Vδ2+ cells did not differ by sex, nor did expression of activation, differentiation, or cytotoxicity markers. These parameters also did not differ significantly between cCMV+ infants who were small for gestational age (SGA) or microcephalic at birth and those who were not. Because infants in this cohort were born in a region of high malaria endemicity, we also assessed associations with histopathologic evidence of placental malaria. However, we found no difference in γδ T subset frequency, expression of proliferation or activation markers, or CD16 expression between placental malaria-exposed and unexposed infants.

In sum, profiling of cord blood γδ T cells from cCMV-infected and uninfected neonates indicates that Vδ1+ and Vγ9-Vδ2+ T cells expand and undergo a high degree of cytotoxic effector differentiation in response to CMV infection *in utero*, and reveals that γδ T cells from cCMV-infected neonates—regardless of subset—exhibit a greater degree of proliferation, activation, and memory differentiation.

### Transcriptional profiling reveals marked cytotoxic effector differentiation of Vδ1+ T cells in cCMV infection

Next, we sought a deeper, unbiased probing of how γδ T cells alter their transcriptional programming in response to congenital CMV infection. For this analysis, we focused on Vδ1+ T cells, which were the most abundant γδ T subset in the cord blood of most infants, particularly those with cCMV. We sorted Vδ1+ T cells (irrespective of γ chain) from the cord blood of 6 cCMV+ and 4 cCMV-neonates and processed them for parallel single-cell RNA- and TCR sequencing to investigate the impact of cCMV infection on Vδ1+ T cell gene expression programs and TCR repertoires.

After processing, sequencing, and quality control, we recovered gene expression data from 11,203 total Vδ1+ T cells from cCMV- individuals and 15,709 total cells from cCMV+ individuals for analysis (average gene number = 675; average Unique Molecular Identifier or UMI number = 1,021). Dimensionality reduction via UMAP and Seurat’s unsupervised Louvain clustering identified seven major clusters among Vδ1+ T cells **(Figure 2A)**. The top ten differentially expressed gene markers for each cluster demonstrated their diverse expression profiles **(Figure 2B)**. The largest cluster (Cluster 0, containing 29.4% of the total cells) was marked by high expression of genes indicative of naïve T cells and factors associated with stemness, such as *TCF7*, *LEF1*, *IL7R,* and *LTB*^41,42^. Cluster 6 broadly shared this transcriptomic profile, with the addition of high *PIK3IP1* expression, a negative regulator of the PI3K-Akt signaling pathway. In contrast, cells in Cluster 2 exhibited highly cytotoxic Type I programming, characterized by the upregulation of multiple granzymes (*GZMH, GZMB, GZMA),* granulysin (*GNLY*), perforin (*PRF1),* and other cytotoxic lymphocyte-associated genes (*NKG7, CD8A)*. Cluster 3 had high expression of amphiregulin, a hallmark wound healing cytokine produced by multiple cell types^43–45^, in addition to transcription factors associated with thymic development (*NR4A1, NR4A3)*^46^. The differentially expressed markers for Cluster 5 contained genes related to innate-like and Type 3 signatures, such as *S100A4, S100A6, KLRB1,* and *ZBTB16* (which encodes the transcription factor PLZF)^39^. To further substantiate the functional identities of these clusters, we scored the clusters using γδ T cell gene modules derived from the literature **(Supplementary Table 3)**^39^. Using the built-in Seurat function FindModuleScore, the average expression levels for three gene sets—naïve γδ T, Vδ1+ cytotoxic lymphocyte, and Type 3 immunity—were calculated for each cell. The distribution and median of scores for each cluster **(Figure 2C)** supported our annotations based on top DEGs: Clusters 0 and 6 scored the highest out of all the clusters for the naïve gene set, whereas Cluster 2 scored both highest for the Vδ1 cytotoxic lymphocyte set and lowest for the naïve set, and Cluster 5 scored the highest for Type 3 immunity. Thus, gene expression profiling indicates that cord blood Vδ1+ T cells are transcriptionally and functionally diverse.

**Figure 2.**
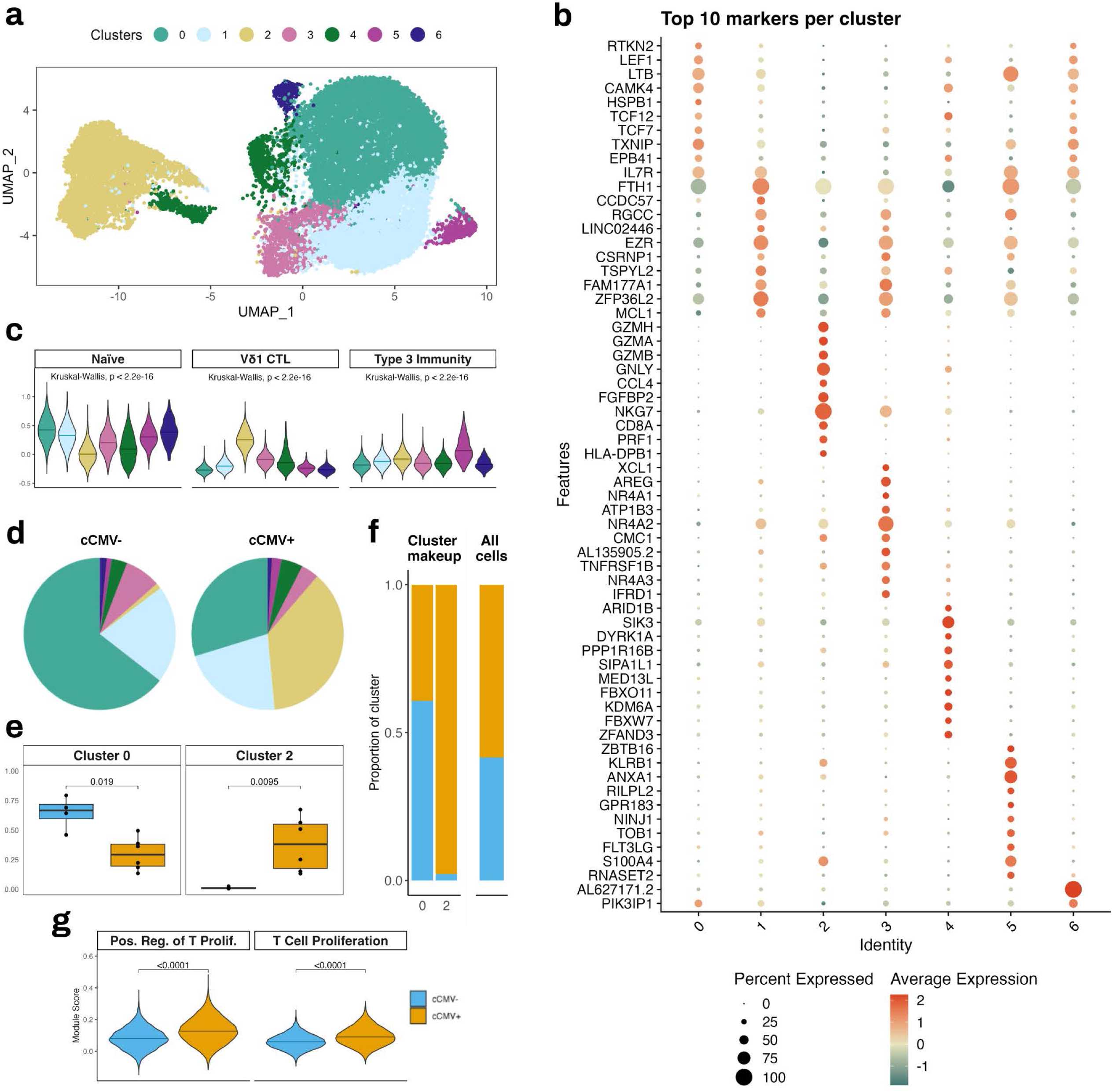
γδ T cells from cCMV+ neonates exhibit highly cytotoxic effector programming and greater proliferation. **A)** UMAP projection of sorted Vδ1+ γδ T cells from cord blood of cCMV- (n = 4) or cCMV+ newborns (n = 6). **B)** Dotplot of the top 10 gene markers per cluster. **C)** Violin plot of module score per cluster for naïve gene set, cytotoxic Vδ1 gene set, or Type 3 immunity gene set. Horizontal bars indicate median scores for all cells in the cluster. P-values were determined by Kruskal-Wallis test. **D)** Pie graphs displaying the cluster distribution for cells from cCMV- individuals (n = 11,203) or all cCMV+ individuals (n = 15,709). **E)** Boxplot displaying frequency of cells that appear in Cluster 0 or Cluster 2 per individual. Medians and IQR for cCMV- (blue) or cCMV+ (orange) neonates shown. **F)** Stacked barplots showing the makeups of Cluster 0 or Cluster 2 (the proportion that is cCMV- vs. cCMV+ cells), left, and the makeup of the entire object, right. **G)** Violin plot of module scores for cell proliferation gene sets from MSigDB for cCMV- cells (blue) or cCMV+ cells (orange); Positive Regulation of T Cell Proliferation gene set (GO:0042102), T Cell Proliferation (GO:0042098). For *E* and *G:* p-values were determined by Wilcoxon rank sum test.

Next, we assessed how the distribution of Vδ1+ T cells among these clusters differed between cCMV-infected and uninfected neonates **(Figure 2D)**. We found that infants infected *in utero* with CMV had a dramatically larger proportion of cells in the cytotoxic Cluster 2 and substantially fewer cells in the naive Cluster 0 compared to uninfected neonates **(Figure 2D)**. This lower proportion of naïve cells in cCMV+ infants corroborates our observations by flow cytometry **(Figure 1D)**. These differences were consistently observed across individuals, with a significantly lower frequency of naïve Cluster 0 cells and a significantly higher frequency of cytotoxic Cluster 2 cells amongst cCMV+ infants **(Figure 2E)**. Strikingly, nearly all cells in Cluster 2 (97.8%) originated from cCMV+ infants **(Figure 2F)**, despite the roughly equal total cell numbers between groups overall. Cells in Clusters 4 and 5 were also disproportionately derived from cCMV+ infants **(Supplementary Figure 2A)**; however, the overall frequencies of cells in these clusters were lower and more variable **(Supplementary Figure 2B)**. The naïve Cluster 0, in contrast, had a greater proportion of Vδ1+ cells derived from cCMV-uninfected infants **(Figure 2F)**, as did Clusters 3 and 6 **(Supplementary Figure 2A)**.

Finally, having observed by flow cytometry that γδ T cells of all subsets were more proliferative in cCMV+ infants **(Figure 1)**, we were interested in whether γδ T cells from cCMV+ infants generally possessed a more broadly proliferative transcriptional program compared to cCMV-infants. We therefore scored all cells using the gene sets that comprise the Gene Ontology terms “Positive Regulation of T Cell Proliferation” and “T Cell Proliferation” from the Molecular Signatures Database^47^. Cells from cCMV+ neonates scored higher for both proliferation-associated gene modules than cells from cCMV-neonates **(Figure 2G)**. These findings together suggest that highly cytotoxic Vδ1+ T cell effectors proliferate and expand to mediate an antiviral response to CMV *in utero*.

### TCR repertoires of cCMV+ infants display oligoclonal γδTCR expansions

We additionally sequenced the TCRs from these same Vδ1+ cells to determine how cCMV infection alters their diversity and clonal composition and to explore their developmental origin. We recovered TCR sequences for 18,619 cells (69.5% of total cells), identifying a TCRδ in 79% of cells and a TCRγ in 74.7%. To visualize the diversity of the TCRγ and TCRδ repertoire within each individual and the degree of clonotype sharing across individuals, we generated treemap plots, with each TCR clonotype colored by the number of other individuals also possessing that clonotype **(Supplementary Figure 3)**. These plots illustrate that all cCMV+ neonates exhibited oligoclonal expansions, with some clonotypes comprising up to 14% of total Vδ1+ cells in the infant’s repertoire. Such oligoclonal expansions were not observed in uninfected neonates.

We compared the overall repertoire diversity by calculating the proportion of each individual’s repertoire that was composed of a unique CDR3δ or CDR3γ sequence found on only one cell. While unique sequences comprised 80-90% of the repertoire in uninfected neonates, among cCMV-infected neonates, unique CDR3δ and CDR3γ sequences comprised a significantly smaller proportion of the total repertoire **(Figure 3A)**, indicating the presence of expanded clonotypes within these individuals. These oligoclonal expansions are further evident when examining the average cumulative proportion of the top ten sequences for the TCRδ and TCRγ chains. The additive proportion of the repertoire was much higher for each clone in cCMV+ individuals compared to cCMV-individuals **(Figure 3B)**.

**Figure 3.**
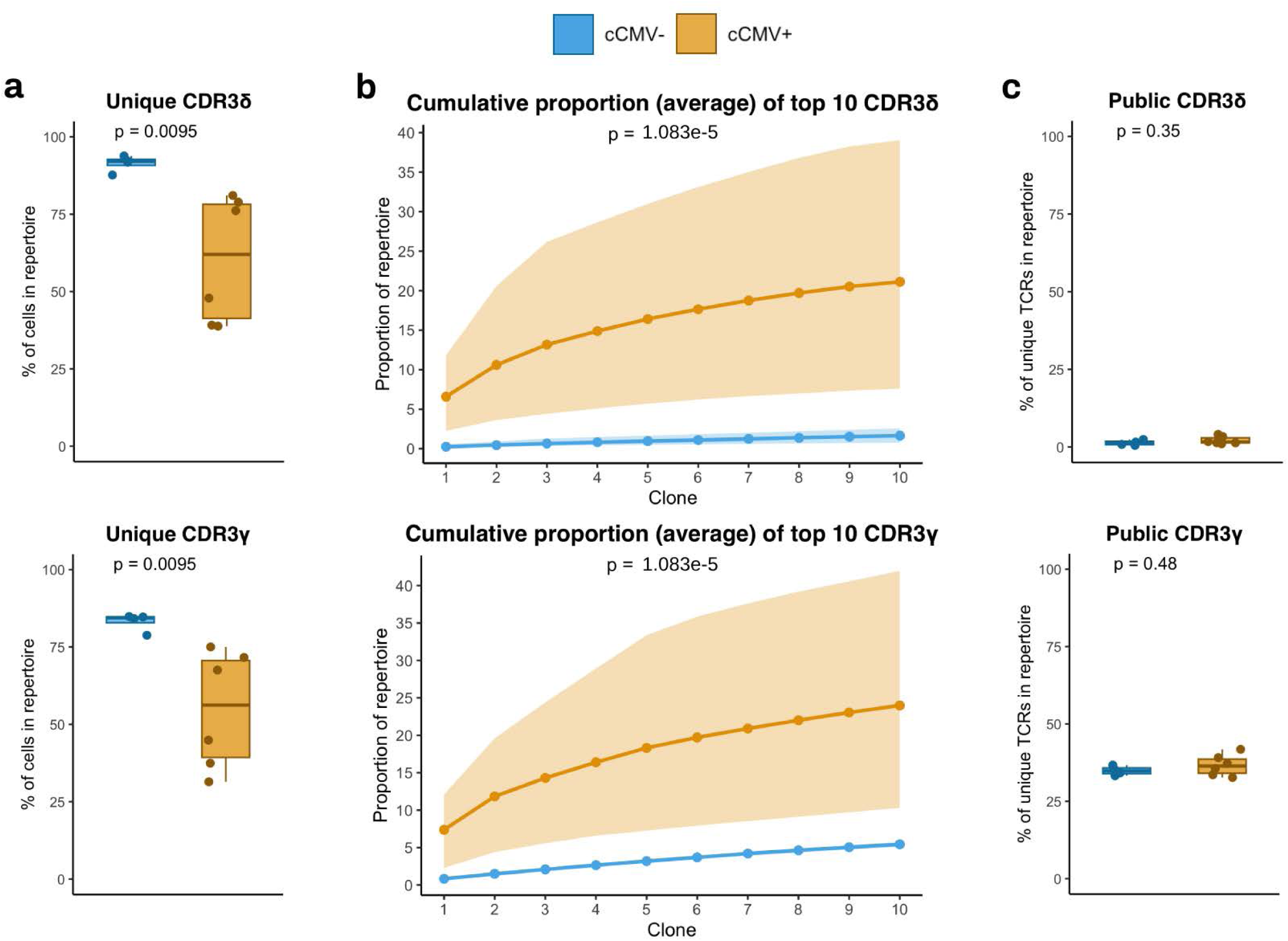
γδTCR repertoires of cCMV+ infants are more oligoclonal than those of cCMV-infants and possess clonotypic expansions. **A)** Percentage of total cells per an individual’s repertoire that have a TCRδ (top) or TCRγ (bottom) that appears on only one cell, for cCMV- (blue) or cCMV+ (orange). Each point is an individual subject. **B)** Accumulation frequency plots representing the cumulative percentage of the repertoire for the top 10 most prevalent TCRδ (top) or TCRγ (bottom) clonotypes in cCMV-individuals (blue) or cCMV+ individuals (orange). Points and solid line indicate mean for the group while shaded area indicates the minimum and maximum values. P-value was determined by Kolmogorov-Smirnov two-sample test. **C)** Percentage of distinct TCRδ (top) or TCRγ (bottom) sequences that are public per an individual’s repertoire for cCMV- (blue) or cCMV+ (orange). Each point is an individual subject and clonotypes are defined using the amino acid sequence for the TCRδ or TCRγ chain. For *A* and *C:* statistical testing was done via Wilcoxon rank sum test.

We also assessed the degree to which γδTCR sequences were “public,” or shared across individuals in this study. On average, the publicity of TCRγ was higher than TCRδ, consistent with prior reports. There was no difference in the overall proportion of public TCRs between cCMV-infected and uninfected infants for either TCRδ or TCRγ **(Figure 3C)**. However, there was a striking tendency for the TCRs that were clonally expanded to also be public **(Supplementary Figure 3)**, perhaps suggesting the existence of highly conserved TCR sequences that can bind viral or host-encoded ligands in the setting of cCMV.

### Vδ1+ T cells from cCMV+ individuals are enriched for fetal-associated TCR characteristics

Prior studies have shown that γδ T cells that arise during early fetal development share a set of distinct TCR features that distinguishes them from those that predominate in late gestation and postnatally. These features include a low number of N additions (random nucleotide additions within the gene junctions), shorter CDR3 lengths, and preferential use of particular joining (J) genes in the TCRδ^48^. Because congenital CMV infection often occurs during a window in which these early progenitors predominate, we investigated whether congenital infection results in a persistent expansion of γδ T cells from this early fetal wave and whether cells deriving from these progenitors retain distinct programming. To investigate this, we first compared the overall γδTCR repertoires of cCMV-infected and uninfected infants for the presence of features associated with γδ T cells generated during an early fetal wave.

We found that TCRs of cCMV+ infants had significantly shorter median CDR3δ lengths than those of cCMV-uninfected infants **(Figure 4A)**. In addition, the distributions of the number of N additions in the TCRδ and TCRγ were significantly left-shifted in cCMV+ neonates compared to uninfected neonates, indicating that there are fewer N additions within both these TCR chains **(Figure 4B)**. Specifically, cCMV+ infants were noted to have an enrichment of cells with zero TCRδ N additions (2.78% vs. 0.40% in cCMV-). Compared to TCRδ, the overall distribution of TCRγ N additions appeared to be less impacted by cCMV; however, there was still a higher proportion of cells with zero N additions in cCMV-infected neonates (11.5% vs. 8.43%) **(Figure 4B)**. The enrichment of cells with zero and few N additions was further confirmed by calculations of the mean number of TCRδ N additions per individual; cCMV+ neonates had significantly lower means compared to cCMV- **(Figure 4C)**. Mean TCRγ N additions also trended lower in cCMV+ neonates **(Figure 4C)**.

**Figure 4.**
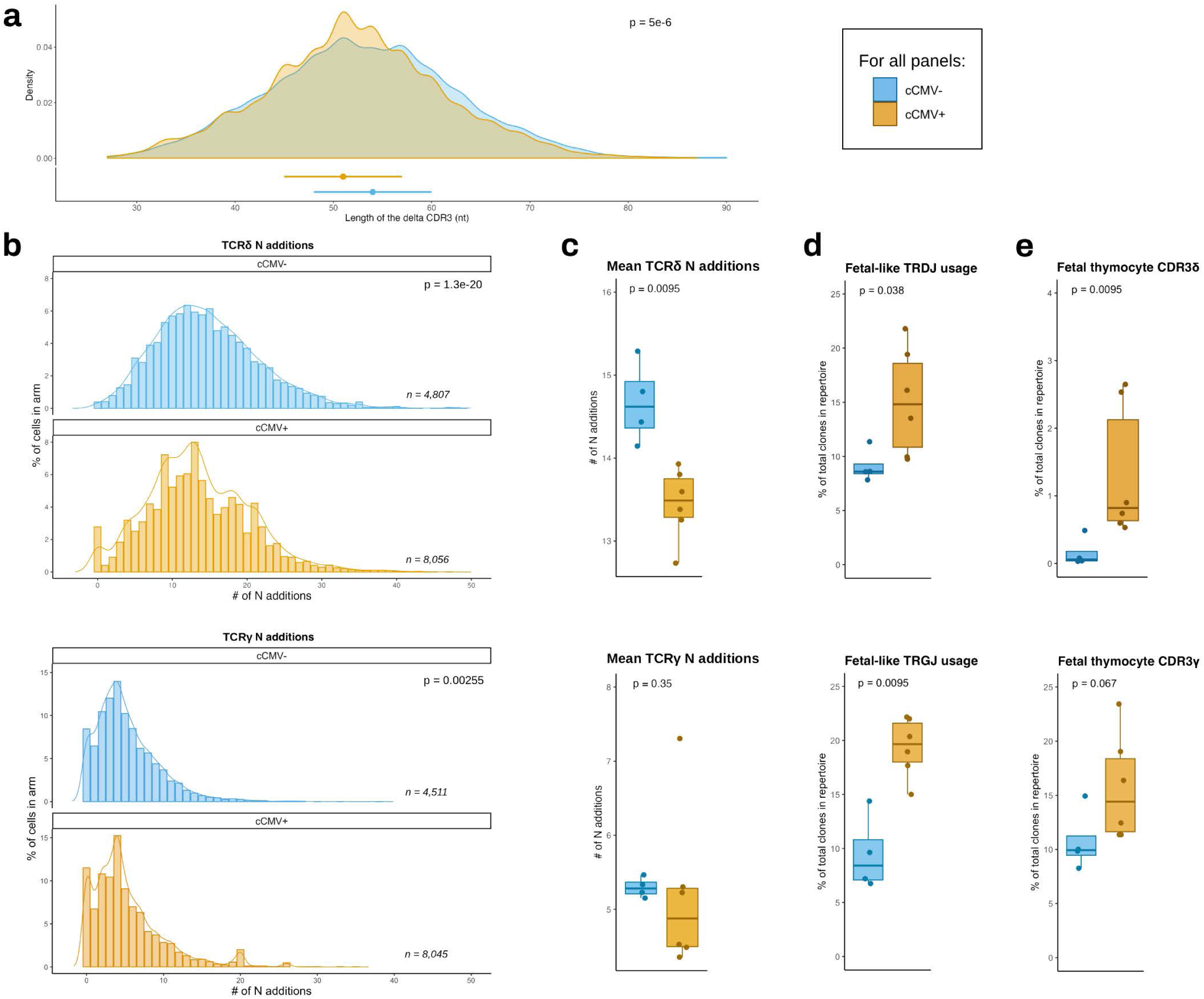
γδTCRs in cCMV+ neonates are enriched for fetal-associated features. **A)** Density histograms of the length in nucleotides of the CDR3δ for cells from cCMV- or cCMV+ neonates. Point interval plots below show median and IQR per group. **B)** Histograms with density overlay of the number of N additions in TCRδ (top) and TCRγ (bottom) for cells from cCMV- or cCMV+ neonates. **C)** Mean number of N additions per individual in TCRδ (top) and TCRγ (bottom) for cells from cCMV- or cCMV+ neonates. **D)** Percentage of cells with fetal-like *TRDJ* (top) or *TRGJ* (bottom) usage for cells from cCMV-or cCMV+ neonates. **E)** Percentage of cells with CDR3δ (top) or CDR3γ (bottom) sequences found in 24-25 gestational week fetal thymi for cells from cCMV- or cCMV+ neonates. For all panels, color corresponds to cCMV status (cCMV- = blue; cCMV+ = orange) and p-values were determined by Wilcoxon rank sum test. For TCRδ: cCMV- n = 4,807, cCMV+ n = 8,065. For TCRγ: cCMV- n = 4,551, cCMV+ n = 8,045. For *C-E:* Median and IQR shown, each point is an individual.

We also analyzed *TRDJ* usage, as it has been shown that fetal thymocytes preferentially use *TRDJ2* or *TRDJ3*, whereas the vast majority of postnatal thymocytes use *TRDJ1*^34^. We found that cCMV-infected neonates had a higher percentage of γδ T cells with fetal-like *TRDJ2* and *TRDJ3* usage **(Figure 4D)**. We observed a similar trend for *TRGJ* usage, with a higher frequency of cells expressing *TRGJP1* (the *TRGJ* gene preferentially used in early fetal-origin cells) in infants with cCMV **(Figure 4D)**^34^. Lastly, we investigated whether CDR3 sequences that we observed in cCMV+ and cCMV- infants matched previously reported γδTCR sequences from human fetal thymocytes at mid-gestation. Using publicly available single-cell TCR repertoires from 6 fetal thymi (estimated gestational ages from 14 to 22 weeks^49^), we observed that for both CDR3δ and CDR3γ, cCMV+ infants had a higher proportion of CDR3 sequences matching those present in fetal thymocytes compared to cCMV- infants **(Figure 4E).**

Together, examination of the TCR repertoire revealed that infection with CMV *in utero* results in a significant enrichment of γδ T cells possessing characteristics associated with early fetal development, suggesting that cCMV infection leads to an expansion of cells derived from early progenitors that is maintained until birth.

### Clonally expanded γδ T cells from cCMV+ infants are cytotoxic and frequently possess fetal-like γδTCRs

Next, we focused on clonally expanded Vδ1+ T cells within cCMV+ individuals to characterize their transcriptional programming and TCR characteristics. The single-cell immune profiling package scRepertoire was used to calculate the frequency of each TCR within an individual infant’s repertoire and categorize them as rare (<0.1% of the repertoire), expanded (0.1-1%), or hyperexpanded (>1%)^50^. Confirming what was observed from the treemap plots and computed accumulation frequencies **(Figures S3 and 3B)**, the repertoires of cCMV+ infants included a greater frequency of γδTCR clonotypes that were defined as “expanded” compared to cCMV- infants. Furthermore, only cCMV+ individuals had “hyperexpanded” clonotypes **(Figure 5A)**. This was true for both the CDR3δ and CDR3γ. Of note, one of the expanded clones matched a previously described public fetal clone (δ1- CALGELGDDKLIF/ γ8 - CATWDTTGWFKIF) that was detected in nearly all of the cCMV+ neonates in a European study^27^. In our cohort, this germline-encoded CDR3δ was present in 5 of 6 infants with congenital CMV but was absent in all 4 cCMV- neonates (**Supplementary Table 5**). Additional expanded CDR3δ clonotypes were observed, several of which were present in more than one cCMV+ infant, although some of these clonotypes were present in uninfected infants as well. The expanded CDR3δ clonotypes paired with a wide variety of TRVG genes (**Supplementary Table 5**), including *TRGV9*, which is notable, as it was previously reported that expansion of γδ T cells in CMV-infected newborns is restricted to Vγ9- cells^27^. This broad Vγ chain usage suggests that many Vδ1+ T cells, and not just the previously reported Vγ8+Vδ1+ cells, may respond to CMV infection *in utero*, although the reactivity of these novel clonotypes to CMV-infected cells remains to be confirmed.

**Figure 5.**
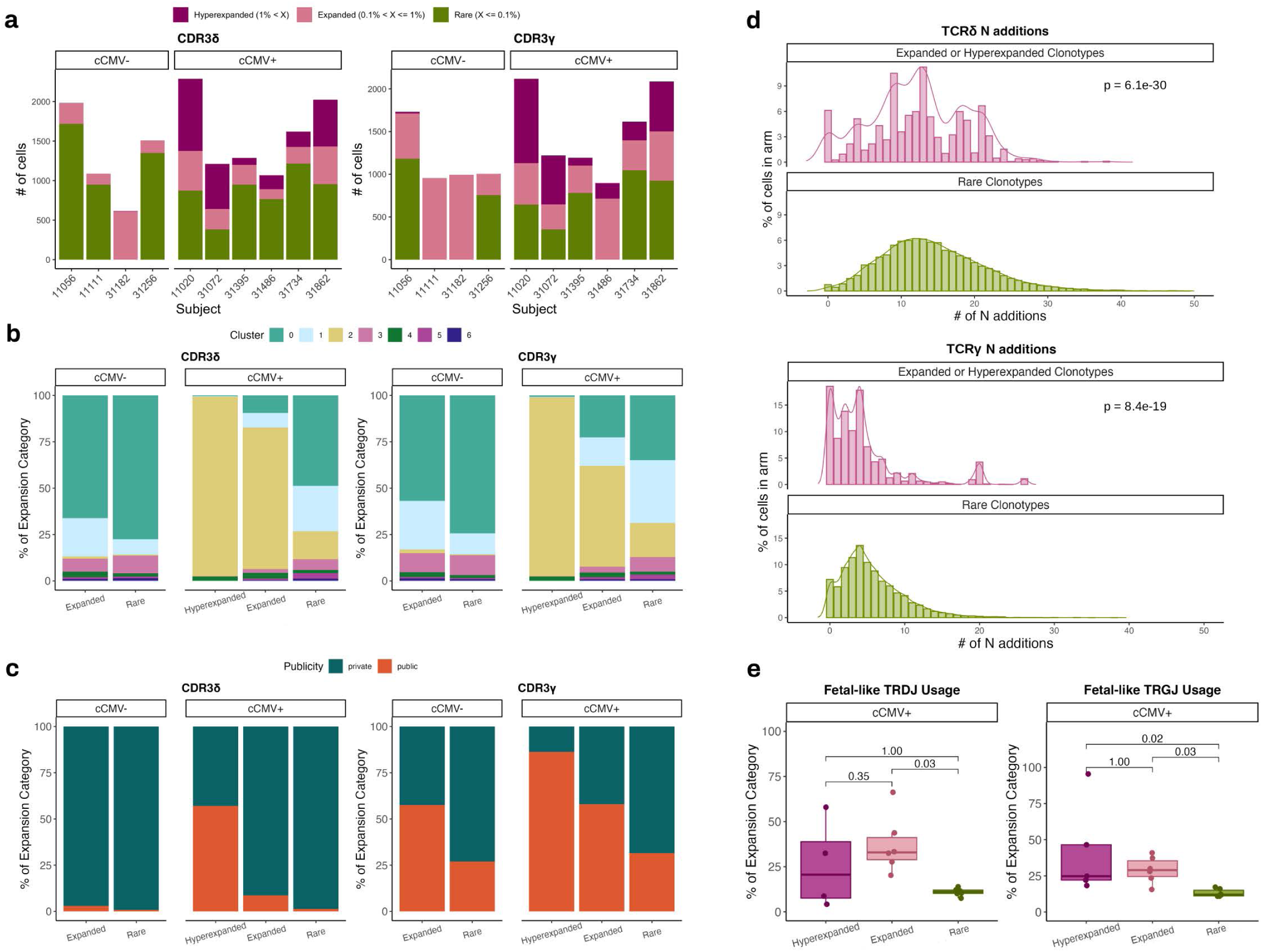
Clonally expanded γδTCRs in cCMV+ neonates have cytotoxic programming and fetal-derived characteristics. **A)** Number of cells in hyperexpanded (1% < X), expanded (0.1% < X <= 1%), or rare (X <= 0.1%) clonal expansion categories per cCMV- or cCMV+ individual. Expansion was calculated within an individual’s repertoire. **B)** Proportion of cells in each cluster per clonal expansion category for cells from cCMV- or cCMV+ infants. **C)** Proportion of cells in each clonal expansion category that are public versus private for cells from cCMV- or cCMV+ infants. For *A-C:* CDR3δ left, CDR3γ right; *hyperexpanded*: cCMV+ n = 2,124; *expanded*: cCMV- n = 463, cCMV+ n = 1,820, *rare*: cCMV- n = 6,734, cCMV+ n = 7,574. **D)** Top: Histograms with density overlay of the # of N additions in TCRδ for cells in expanded or hyperexpanded clonal expansion categories (n = 3,420) or rare category (n = 9,031). Bottom: Histograms with density overlay of the # of N additions in TCRγ for cells in expanded or hyperexpanded clonal expansion categories (n = 3,543) or rare category (n = 9,053). P-values were determined by Wilcoxon rank sum test. **E)** For cCMV+ individuals, the percentage of cells within each clonal expansion category with fetal-like *TRDJ* usage (left) and fetal-like *TRGJ* usage (right). Each point is an individual. P-values were determined by a Kruskal-Wallis with Dunn’s post-hoc test and Bonferroni correction.

Having defined these expansion categories, we sought to determine whether there was common functional programming between the expanded clones and whether this was influenced by *in utero* CMV exposure. We leveraged paired single-cell sequencing to assess the relationship between transcriptional programming and clonal expansion. We observed that among cCMV-uninfected neonates, most cells, whether from rare or expanded clonotypes, clustered with naïve cells (Cluster 0). In contrast, among cCMV+ infants, clones that were expanded and hyperexpanded were predominantly in the highly cytotoxic Cluster 2, whereas rare clones were more likely found in the naïve Cluster 0 and Cluster 1 **(Figure 5B)**. In addition, cells with expanded or hyperexpanded CDR3δ were more likely to be public than rare clonotypes, specifically in cCMV-infected neonates, but not in cCMV-uninfected neonates **(Figure 5C)**. Interestingly, this was not the case with CDR3γ, for which expanded clonotypes were more often public regardless of cCMV exposure. This is likely due to the less diverse nature of CDR3γ sequences at baseline, whereas CDR3δ sequences tend to be more variable.

We next assessed whether the clonally expanded Vδ1 cells possessed TCR features indicative of early fetal origin. Comparison of the distributions of TCRδ or TCRγ N additions between expanded and hyperexpanded clonotypes versus rare clonotypes in cCMV+ individuals showed that indeed the expanded and hyperexpanded clonotypes were enriched for CDR3 sequences with few or no N additions (**Figure 5D)**. The expanded clones also had a bias toward usage of the fetal-like *TRDJ2* or *TRDJ3* genes as well as the fetal-like *TRGJP1* **(Figure 5E**), further suggesting that they disproportionately derive from an early wave of fetal development.

In conclusion, combined scRNAseq and scTCRseq revealed an oligoclonal expansion of highly cytotoxic effector γδ T cells in the cord blood of neonates driven by *in utero* infection with CMV. We additionally found an enrichment for characteristic fetal γδTCR features that suggest these expanded cells preferentially derive from early progenitors.

### Fetal-like γδ T cells display a cytotoxic, NK-like gene expression program

Finally, we sought to obtain a broader, unbiased view of the transcriptional differences between fetal-like and adult-like γδ T cells found in cord blood. Based on prior studies demonstrating the distinct γδTCR characteristics of fetal thymocytes^34^, we reasoned that cells having 0-4 N additions in the TCRδ were highly likely to originate from early fetal progenitors. We compared gene expression of these cells to that of cells with N additions above the median (>12), which are unlikely to arise from fetal progenitors. Unbiased clustering of γδTCRs from neonatal blood identified low/no N additions as characteristic of fetal-like γδ T clusters^40^; moreover, in explorations of our data using either the KAy-means for MIxed LArge data (KAMILA) clustering algorithm^51^ or the built-in Seurat sub-clustering function, the low versus high N addition dichotomy was consistently a variable that distinguished clusters. Thus, using this framework, we performed differential gene expression on fetal-like versus adult-like Vδ1+ T cells in our dataset.

Fetal-like cells had higher expression of cytotoxic, natural killer (NK)-associated genes like *NKG7*, *GZMB*, *PRF1*, and *TYROBP*, consistent with a pre-programmed effector bias^1,27,30,34^ **(Figure 6A)**. In addition, we observed downregulation of *TGFBR2* and *SMC4*, both of which promote TGF-β signaling^52,53^. TGF-β inhibits T cell differentiation into effectors^54,55^; therefore, inhibition of this pathway would, in turn, lead to enhanced effector differentiation. Gene Ontology enrichment and subsequent network analyses elucidated and confirmed the overall transcriptional programs possessed by fetal-like γδ T cells. As expected from the listed upregulated genes, enriched ontology terms included “NK-mediated killing” and “NK-mediated cytotoxicity” **(Figure 6B)**, while downregulated terms included both those related to differentiation (lymphocyte and T cell differentiation) and those related to regulation of function (“regulation of T cell activation”, “antigen receptor-mediated signaling pathway”) **(Figure 6C)**.

**Figure 6.**
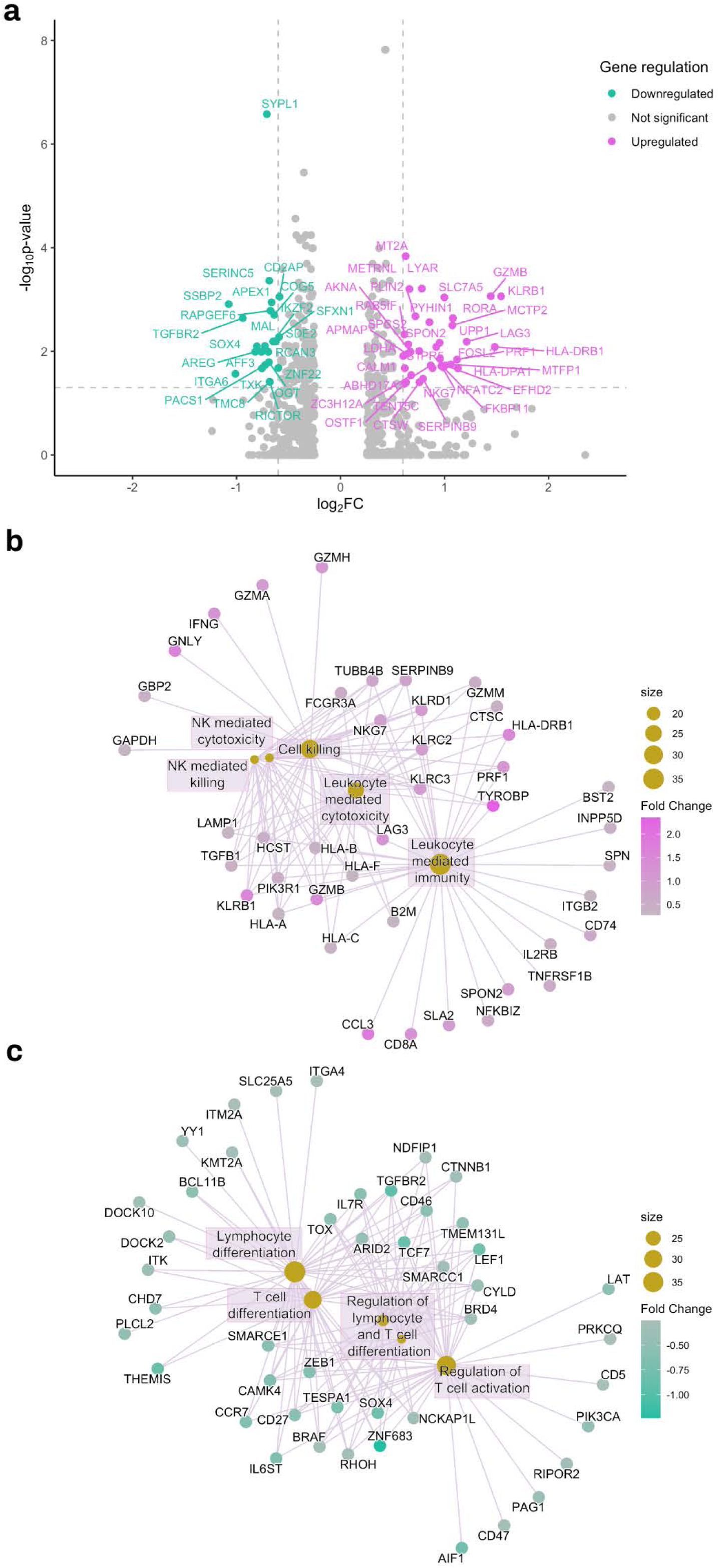
Fetal-derived γδ T cells display NK cell-like transcriptional programming. **A)** Volcano plot depicting genes significantly upregulated (magenta) or downregulated (teal) in fetal-like Vδ1+ T cells versus adult-like Vδ1+ T cells. Horizontal dashed line: adjusted p-value = 0.05; Vertical dashed lines: fold change = −1.5-fold or 1.5-fold. **B)** Gene-concept network plot depicting the top 5 enriched Gene Ontology terms for genes upregulated in fetal-like versus adult-like Vδ1+ T cells. **C)** Gene-concept network plot depicting the top 5 enriched Gene Ontology terms for genes downregulated in like versus adult-like Vδ1+ γδ T cells. *For B and:* Nodes are the enriched Gene Ontology terms and the termini the associated DEGs. Size of the node indicates how many genes were found in that GO term.

In sum, unbiased differential gene expression analysis comparing cord blood γδ T cells possessing fetal-like versus adult-like TCRs demonstrated that fetal-like cells are marked by an NK-like transcriptional program that equips them for cytotoxic effector function *in utero*.

## DISCUSSION

Human antiviral immune defenses evolve rapidly during fetal development and the first years of life in ways that are critically important for viral containment. As the first T cells to develop during human gestation, γδ T cells are on the front line of fetal host defense. However, few studies have examined how fetal γδ T cells respond to a perinatal pathogen. In this report, we found that CMV infection *in utero* drives an oligoclonal expansion of γδ T cells that adopt a highly cytotoxic effector differentiation profile and are enriched for TCR features that suggest they are derived from an early fetal progenitor population.

We found that the increased frequency of γδ T cells in cCMV+ individuals was due in part to oligoclonal expansion of particular Vδ1+ clones that are highly effector-differentiated and predominantly exhibit cytotoxic effector programming. Interestingly, and in contrast to adult CMV infection^28,22^, the majority of these expanded clonotypes were public, or shared among multiple infants. A prior study identified a CMV-reactive public Vγ8+Vδ1+ clone that was shown via spectratyping to be present in the cord blood of five of the six cCMV+ infants studied, but was absent in the circulating repertoire of CMV-uninfected infants at birth^27^. Remarkably, a subsequent analysis of fetal thymocytes noted this germline-encoded TCR sequence to be present in fetal thymocytes at mid-gestation^34^. That expansion of this same γδTCR is seen in Ugandan infants from a distinct geographic area implies that this is a widely conserved response to CMV. In addition to this previously identified δ1-CALGELGDDKLIF/γ8-CATWDTTGWFKIF clonotype, we found several other clonotypes expanded in neonates with cCMV, many of them public. Given the long-term co-evolution of CMV and humans, this raises the intriguing prospect that these TCR sequences have been evolutionarily conserved to mediate antiviral CMV effector functions *in utero*. Further structural analysis of these γδTCRs may identify potential commonalities in structure or biochemical properties, laying the groundwork for identification of viral or host-encoded Vδ1 TCR ligands, which thus far remain largely unknown^23,25^.

Notably, we found that many of the expanded γδ T clonotypes possessed shared TCR features that are suggestive of early fetal origin, including a low number of random nucleotide (N) additions^34^. The TCR repertoire of fetal T cells (both αβ and γδ) is less diverse and more broadly reactive than that of adults^56^. Elegant studies have recently shown that the low junctional diversity seen in γδ T cells that derive from early progenitor populations is due to the low activity of TdT (terminal deoxynucleotidyl transferase), the enzyme that catalyzes the random insertion of N nucleotides^34^. Fetal hematopoietic stem and precursor cells (HSPCs) have intrinsically high expression of the RNA-binding protein Lin28b, which suppresses TdT expression. The resulting low diversity γδTCR repertoire may reflect the distinct priorities of host defense in early life, as both B and T cells arising during early development appear to favor broad reactivity over specificity and rapid effector action over memory formation^56^. Intriguingly, Lin28b may act broadly as a developmental switch between the fetal and adult “layers” of the immune system, as it suppresses the *let-7* family of microRNAs, which influences the transcriptional shift from the effector-biased expression program of early life toward the generation of immune memory in later life^34,56^.

By leveraging paired transcriptomic and TCR sequencing of individual γδ T cells to identify those with TCRs characteristic of fetal progenitor origin, we were able to discern differences in the global gene expression programs of γδ T cells arising from this early wave of development. Overall, we found that these early fetal-origin γδ T cells are poised for rapid effector function and possess transcriptional programs that are heavily NK-like. The resemblance of γδ T cells to NK cells resides not only in their highly cytotoxic programming and expression of cytotoxic granule-associated molecules, but other potential functions as well. In particular, we observed that γδ T cells in cCMV+ neonates upregulate expression of the Fc receptor CD16. We reported in a previous study that the frequency of CD56^neg^CD16^+^ NK cells is increased in neonates with cCMV^57^. Thus, the developing fetus is armed with two populations of Fc receptor-expressing lymphocytes that can leverage transplacentally acquired maternal antibodies for antibody-dependent cellular cytotoxicity. In line with this model, recent studies have shown that high avidity maternal CMV-specific antibodies that are able to engage FcR-mediated cellular immune responses correlate with protection against congenital CMV transmission^58,59^. The ability to harness the specificity of maternal IgG via Fc-mediated immune functions may provide a critical first line of defense against viral infection at a time when the fetus lacks acquired immune memory. This collaboration with maternal-origin antibodies may be one of the many tools evolved by γδ T cells to bridge innate and adaptive immunity in the fetus.

An important limitation of our study is that the timing of mother-to-infant CMV transmission likely varied among participants and could not be determined, as vertical transmission is a clinically silent event. Because human gestation is a period of rapid immune development, the antiviral response of the fetus may differ substantially between early and late gestation. Severe sequelae of cCMV, such as microcephaly, are more likely to occur in infants infected during the first trimester^60–63^ and currently, no therapies are available to mitigate the devastating effects of fetal infection, even when it is identified early. The ability of γδ T cells to mediate rapid HLA-unrestricted antiviral effector functions raises the intriguing prospect of incorporating these broadly CMV-reactive γδTCRs into cellular therapies that could be administered to the fetus *in utero*. Moreover, because γδ T cells do not require HLA for antigen presentation, they are not susceptible to the CMV evasion strategy of HLA downregulation^64^. Although we did not observe a correlation between γδ T cell frequencies and clinical sequelae at birth, it would be intriguing to examine this relationship in a larger cohort of cCMV+ infants. While cCMV infection has devastating consequences for some infants, the majority remain asymptomatic^61,63^, and it is possible that γδ T cells contribute to a partially protective immune response that enables most infants to evade sequelae. Supporting this, murine models have shown that in circumstances where conventional αβ T cells are impaired or absent, γδ T cells alone are able to directly restrict CMV replication and subsequently minimize pathology^17,65^.

In summary, we have demonstrated that γδ T cells expand in response to CMV infection *in utero* and exhibit a high degree of proliferation, activation, and memory differentiation. This expansion is comprised of highly cytotoxic, oligoclonally expanded cells that frequently possess public, fetal-like TCRs. These findings further our understanding of the role of γδ T cells in early life ontogeny and their development and differentiation in response to infection *in utero*. As rapid, broadly reactive, and innate-like effectors, γδ T cells may provide a critical defense during fetal and early life, when adaptive immune responses are lacking.

## METHODS

### Study populations and samples

Cord blood mononuclear cells (CBMCs) were obtained from a subset of infants (n = 85) enrolled in a clinical trial of prenatal and early childhood malaria chemoprevention conducted in the Tororo and Busia Districts, a highly malaria endemic area in Eastern Uganda (PROMOTE-BC1: NCT02163447: PROMOTE-BC3: NCT02793622). Details of the larger study, including eligibility criteria for enrollment and clinical outcomes, have been previously described in detail^66^. All pregnant women in this trial screened negative for human immunodeficiency virus (HIV) at enrollment. Umbilical cord blood was collected at delivery using cord blood collection kits (Pall Medical) and an aliquot of whole cord blood preserved in RNAlater (ThermoFisher) at −80°C. CBMCs were promptly isolated from the remaining cord blood using density gradient centrifugation (Ficoll-Histopaque; GE Life Sciences) and cryopreserved in liquid nitrogen.

### Ethical approval

Written informed consent was obtained from the mothers of all study participants. The study protocol was approved by protocol was approved by Makerere University School of Biomedical Sciences Ethics Committee, Uganda National Council of Science and Technology, and the University of California, San Francisco Research Ethics Committee.

### Congenital CMV screening

To identify congenitally CMV-infected newborns, DNA was extracted from whole cord blood preserved in RNAlater using the QIAamp DNA Blood Mini Kit (Qiagen) according to the manufacturer’s instructions. Extracted DNA was then quantified with a Qubit 4 Fluorometer (Invitrogen™) before screening for CMV viral DNA by quantitative real-time PCR (qRT-PCR) using PowerUp™ SYBR™ Green Master Mix (Applied Biosystems™) on the ABI StepOne Plus system. All samples were screened in duplicate using primers targeting UL123 **(Supplementary Table 1)**^67^. CMV-positive samples were re-tested with a second qRT-PCR using a UL55 primer set to confirm positivity **(Supplementary Table 1)**.

All discordant qRT-PCR results were re-screened by Droplet Digital PCR (ddPCR) as published previously^68^. Briefly, 5 μL of extracted DNA was added to 10 μL of ddPCR 2X PCR master mix (Quantalife, Applied Biosystems) in addition to 1 μL each of 10 μM forward and reverse primers and 1 μL of 5 μM probe targeting UL55; the same primers were used for qRT-PCR and ddPCR **(Supplementary Table 1)**. The combined DNA and master mix (18 μL) was added to the droplet generator strips, placed in the automated droplet generator, and the resulting emulsion transferred to a 96-well PCR plate. After amplification, the number of positive and negative droplets were enumerated using a BioRad QX300 droplet reader based on fluorescence.

### Flow cytometry

CBMCs were thawed, counted, evaluated for viability, and stained for extra- and intracellular targets with antibodies listed in **Supplementary Table 2**. CBMCs were stained with LIVE/DEAD Fixable Near-IR (ThermoFisher) to exclude dead cells. For intracellular staining, CBMCs were fixed using a Cytofix/Cytoperm kit (BD) and stained in Permeabilization/Wash buffer (BD) per manufacturer’s instructions. Data was acquired on an LSR II (BD) flow cytometer using FACS DIVA software.

Compensation was performed using single-color stained UltraComp beads (Invitrogen) and a minimum of 500,000 events were recorded from each sample. Flow cytometry data were analyzed using FlowJo version 10.10.0 (Tree Star). Fluorescence minus one (FMO) controls were used to set gates for all markers. γδ T cells were defined as LIVE/DEAD-negative single cells that were CD14-/CD19-/CD3+ and positive for at least one of the γδTCR chains (Vδ1, Vδ2, or Vγ9) (**Supplementary Figure 1A)**.

### Single-cell RNA and TCR sequencing

#### Library generation and sequencing

For single-cell RNA sequencing (scRNAseq) and single-cell TCR sequencing (scTCRseq), 10,000 Vδ1+ T cells were FACS sorted from cryopreserved CBMCs from four cCMV- neonates and six cCMV+ neonates. Gene expression libraries were constructed using the Chromium Next GEM Single Cell 5’ Library & Gel Bead Kit v1.1 (10X Genomics). TCR libraries were constructed using the Chromium Single Cell V(D)J Enrichment Kit, Human T Cell v1.1 (10X Genomics). Manufacturer’s protocols for library construction were followed, except for the use of custom primers targeting the delta chain of the TCR (TCRδ) and the gamma chain (TCRγ) for TCR enrichment, published previously (**Supplementary Table 1)**^49^. Throughout library construction, Agilent Bioanalyzer High Sensitivity DNA chips and a Bioanalyzer 2100 machine were used to check cDNA and library quality (Agilent Technologies). scRNAseq libraries were pooled for a targeted read depth of 20,000 reads/cell pooled for a targeted read depth of 5,000 reads/cell. Pooled libraries were sequenced on the Illumina NovaSeq6000 platform.

#### Single-cell RNA gene expression demultiplexing, doublet removal, and normalization

After sequencing, Cell Ranger v7.1.0 with the default parameters was used to align the scRNAseq libraries to the human reference genome GRCh38 (UCSC, CA, USA), filter and count barcodes and UMIs, and generate expression matrices. The pooled libraries comprised nine participants and were demultiplexed by using SNP profiles to match individual cells to the original sample. First, pileup files were generated to summarize the reads at alleles of common SNP sites derived from the 1000 Genomes Consortium^69^. Next, Freemuxlet was used to cluster cells on similar SNP profiles and assign droplet type (singlet, doublet, or ambiguous)^70,71^. These Freemuxlet clusters were then mapped to previously generated bulk RNAseq genotypes of the participants using BCFtools v1.10.2. Barcodes designated as singlets were then carried forward into further processing with DoubletFinder to identify and remove heterotypic intra-sample doublets^72^. For DoubletFinder, the data were transformed using Seurat v5.2.1’s SCTransform v2; principal component analysis (PCA) was performed using npcs = 35, nearest neighbors computed via FindNeighbors(), and the data clustered using FindClusters(). DoubletFinder’s paramSweep_v3() was used to estimate the neighborhood size (pK) per library. An effective heterotypic doublet rate was estimated using the doublet proportion calculated by Freemuxlet divided by (number of samples – 1) to represent all possible intersample doublets. Further specific details on DoubletFinder parameters and implementation are published^73^.

#### Single-cell RNA gene expression integration, filtering, dimensionality reduction, and clustering

Following demultiplexing and singlet identification, Seurat v5.2.1 was used to integrate the normalized data using the Canonical Correlation Analysis method built into the IntegrateData() function. For this, we selected the first 3000 variable features to identify integration anchors in Seurat’s

FindIntegrationAnchors() function and then integrated the normalized data using default inputs. Following integration, low-quality cells were filtered out of the resulting Seurat object. A cell meeting any of the following criteria was removed: <250 or > 2000 detected genes; >= 10% mitochondrial gene percentage, or >1% of reads mapping to hemoglobin genes. Mitochondrial and ribosomal percentages, cell cycle state, and percent of reads mapping to TCR genes were regressed out of the integrated, filtered Seurat object using SCTransform with vst.flavor = “v2”.

For clustering, we performed PCA and selected the first 40 principal components as the input for dimensionality reduction using Uniform Manifold Approximation and Projection (UMAP) with n.neighbors = 20, min.dist = 0.5 (all other parameters were default). The shared nearest-neighbor graph was then constructed using the same principal components and this graph subsequently used to identify cell clusters via Seurat’s FindClusters() k-nearest neighbor clustering with the Louvain algorithm.

Clustering was performed at a series of resolutions, and the package clustree v0.5.1^74^ was used to visualize how cells move as the number of clusters increases, thereby aiding in selecting an appropriate resolution (resolution = 0.2). Marker genes for each cluster were identified using FindAllMarkers() and Wilcoxon rank-sum test method with a logfc.threshold = 0.264 (average fold change of at least 1.2), min.pct = 0.05, and an adjusted p-value of at least 0.05.

#### Module scoring

Scores for gene set modules were calculated using Seurat’s AddModuleScore() function. Naïve, Vδ1 Cytotoxic Lymphocyte, and Type 3 Immunity gene sets derive from published work^39^. All genes associated with the Gene Ontology terms “Positive Regulation of T Cell Proliferation” (GO: 0042102) or “T cell Proliferation” (GO: 0042098) were downloaded from MSigDB^47,75^. See **Supplementary Table 3** for full gene sets.

### Single-cell TCR expression data analysis

#### γδTCR annotation and clonotype calling

The scTCRseq libraries were aligned to the GRCh38 reference genome (UCSC, CA, USA) using Cell Ranger vdj v7.1.0 and v3.0.2. Because Cell Ranger vdj versions >= 3.1 correctly align γδTCR contigs but do not accurately annotate them, the aligned output from Cell Ranger vdj v.7.1.0 was processed via the Python package Dandelion v0.5.2 for re-annotation^76^. Briefly, the FASTA and contig annotation files resulting from Cell Ranger vdj were put through the standard Dandelion pre-processing, which annotates the contigs according to the latest ImMunoGeneTics (IMGT) human reference libraries and outputs AIRR-formatted calls. As a secondary method, scTCRseq libraries were also aligned to the genome using Cell Ranger v3.0.2. From these, we only kept the contig calls for the set of cells that did not already have γδTCR sequences called by v7.1.0 and Dandelion.

To determine TCR clonotypes, we used the Dandelion-annotated sequences as input for Scirpy v0.19.0, first using scirpy.pp.ir_dist() to compute distances between CDR3 nucleotide (nt) or amino acid (aa) sequences. We then defined clonotypes with identical CDR3 sequences at the nucleotide level using scirpy.tl.define_clonotypes() with default parameters and dual_ir = “primary”. We defined clonotypes with identical CDR3 sequences at the amino acid level using scirpy.tl.define_clonotype_clusters() with parameters dual_ir = “primary,” metric = “identity,” and sequence = “aa”. The clonotype calls, gene segment calls, CDR3 sequences, and junction sequences output from Scirpy were then incorporated into the Seurat object as metadata based on cellular barcodes.

#### Calculating N additions

The web-based Junctional Analysis Tool from IMGT was used to calculate N additions (non-templated nucleotides added at TCR gene junctions). For each cell, the V and J gene names and alleles, plus the junction nucleotide sequence, were put into FASTA format and then submitted via the website for analysis. All additional options were left as the defaults. The total number of nucleotides at all N regions was summed to yield the final number of N additions, and the delta and the gamma chains were analyzed separately.

#### scRepertoire, clonal frequency, and publicity

scRepertoire v2.2.1^50^ was used for clonal and repertoire analysis. The clonotype information from scirpy was reformatted to fit scRepertoire’s expected data structure, and the Seurat object split based on participant ID. scRepertoire’s combineExpression() function was subsequently employed to calculate the clonal frequency and proportion for each clonotype within an individual’s TCR repertoire, and then that information was incorporated back into the Seurat object as metadata based on matching cellular barcodes. For the above, a clonotype was defined using the amino acid sequences of either the gamma or delta CDR3. In some cells where sequencing failed to capture information for both chains, the missing chain information was imputed using cells from the same individual that possessed an identical gamma or delta CDR3 and that did have both TCR chains sequenced.

Publicity was defined for the TCRγ and TCRδ chains separately, by comparing the CDR3 amino acid sequence to all other existing CDR3 sequences across all individuals for the respective chain. A CDR3 sequence was considered private if it appeared only in one individual and public if it appeared in two or more, with an additional calculation for how many other individuals it appeared in.

#### Differential gene expression analysis

For differential gene expression analysis between fetal-derived versus postnatal γδ T cells, we used the Model-based Analysis of Single Cell Transcriptomics (MAST) algorithm (https://github.com/RGLab/MAST/)^77^ with cluster identity and cCMV status as latent variables and modified to use particpant as a random effect variable variable to account for autocorrelation of per cell gene expression within a subject, from here (https://github.com/CBMR-Single-Cell-Omics-Platform/SCOPfunctions/blob/main/R/differential_expression.R). MAST was used to determine genes and p-values while the default calculation from Seurat was used to determine average log fold change.

For network plots, the upregulated differentially expressed genes (DEGs) or downregulated DEGs derived via MAST above were used as the gene list for Gene Ontology enrichment in clusterProfiler v.4.14.4. The resulting enriched gene ontology terms and associated genes were then plotted using the cnetplot() function from the package enrichplot to create network plots.

### Data visualization

The R packages ggplot2^78^, dittoSeq^79^, ggridges^80^, and colorspace^81^ were used to plot the flow cytometry and scRNA- and TCRseq data. Treemap plots were made using the package treemapify^82^.

### Statistical analysis

Statistical testing for all plots was done in R v4.4.2 using the package ggpubr v0.6.0^83^. Two group comparisons were performed using an unpaired Wilcoxon rank-sum test; p-values < 0.05 were considered significant. Multiple group comparisons were conducted using Kruskal-Wallis tests, with Dunn’s post-hoc analysis applied to evaluate pairwise differences. Two-group comparisons of cumulative frequency distributions were performed using the Kolmogorov-Smirnov test; p-values < 0.05 were considered significant.

## ACKNOWLEDGMENTS

We are grateful to the study participants and their families who made this work possible. The authors would like to thank Dr. David Vermijlen and his group, especially Dr. Guillem Sánchez-Sánchez, for their insightful suggestions, γδTCR primers, and assistance with the N addition code. The resources provided by UCSF’s Parnassus Flow Core CoLab (RRID: *SCR_018206,* NIH 1S10OD021822-01) and sequencing by the UCSF Center for Advanced Technology (NIH 1S10OD028511-01 grants) were instrumental in obtaining the data for this study.

## AUTHOR CONTRIBUTIONS

**JL**: Conceptualization, Data curation, Analysis, Funding acquisition, Investigation, Software, Visualization, Writing – Original draft preparation; **AVV**: Conceptualization, Analysis; **JT**: Investigation; **BD:** Investigation; **MEO:** Investigation, Software; **PCC:** Investigation; **CBN:** Investigation; **MI:** Investigation; **GF:** Supervision; **RP:** Software, Supervision; **FN:** Investigation; **GD:** Funding acquisition, Resources; **AK:** Resources; **MM:** Resources; **MEF:** Funding acquisition, Resources, Supervision, Writing – Review and editing

## Notes

### Competing Interest Statement

The authors have declared no competing interest.

## References

1. Davenport, M. P., Smith, N. L. & Rudd, B. D. Building a T cell compartment: how immune cell development shapes function. Nat. Rev. Immunol. 20, 499–506 (2020).

2. McVay, L. D., Jaswal, S. S., Kennedy, C., Hayday, A. & Carding, S. R. The generation of human gammadelta T cell repertoires during fetal development. J. Immunol. Baltim. Md 1950 160, 5851–5860 (1998).

3. Morita, C. T., Parker, C. M., Brenner, M. B. & Band, H. TCR usage and functional capabilities of human gamma delta T cells at birth. J. Immunol. Baltim. Md 1950 153, 3979–3988 (1994).

4. Gibbons, D. L., et al. Neonates harbour highly active γδ T cells with selective impairments in preterm infants. Eur. J. Immunol. 39, 1794–1806 (2009).

5. Vantourout, P. & Hayday, A. Six-of-the-best: unique contributions of γδ T cells to immunology. Nat. Rev. Immunol. 13, 88–100 (2013).

6. Goriely, S., et al. A Defect in Nucleosome Remodeling Prevents IL-12(p35) Gene Transcription in Neonatal Dendritic Cells. J. Exp. Med. 199, 1011–1016 (2004).

7. Renneson, J., et al. IL-12 and type I IFN response of neonatal myeloid DC to human CMV infection. Eur. J. Immunol. 39, 2789–2799 (2009).

8. Farrington, L. A., et al. Opsonized antigen activates Vδ2+ T cells via CD16/FCγRIIIa in individuals with chronic malaria exposure. PLOS Pathog. 16, e1008997 (2020).

9. Couzi, L., et al. Antibody-dependent anti-cytomegalovirus activity of human γδ T cells expressing CD16 (FcγRIIIa). Blood 119, 1418–1427 (2012).

10. Brandes, M., Willimann, K. & Moser, B. Professional Antigen-Presentation Function by Human γδ T Cells. Science 309, 264–268 (2005).

11. Brandes, M., et al. Cross-presenting human γδ T cells induce robust CD8+ αβ T cell responses. Proc. Natl. Acad. Sci. 106, 2307–2312 (2009).

12. Ramsburg, E., Tigelaar, R., Craft, J. & Hayday, A. Age-dependent Requirement for γδ T Cells in the Primary but Not Secondary Protective Immune Response against an Intestinal Parasite. J. Exp. Med. 198, 1403–1414 (2003).

13. Hayday, A. C. γδ Cells: A Right Time and a Right Place for a Conserved Third Way of Protection. Annu. Rev. Immunol. 18, 975–1026 (2000).

14. Pinninti, S. & Boppana, S. Congenital Cytomegalovirus Infection Diagnostics and Management. Curr. Opin. Infect. Dis. 35, 436–441 (2022).

15. Manicklal, S., Emery, V. C., Lazzarotto, T., Boppana, S. B. & Gupta, R. K. The “Silent” Global Burden of Congenital Cytomegalovirus. Clin. Microbiol. Rev. 26, 86–102 (2013).

16. Lanzieri, T. M., Dollard, S. C., Bialek, S. R. & Grosse, S. D. Systematic review of the birth prevalence of congenital cytomegalovirus infection in developing countries. Int. J. Infect. Dis. IJID Off. Publ. Int. Soc. Infect. Dis. 22, 44–48 (2014).

17. Khairallah, C., Déchanet-Merville, J. & Capone, M. γδ T Cell-Mediated Immunity to Cytomegalovirus Infection. Front. Immunol. 8, 105 (2017).

18. Déchanet, J., et al. Implication of γδ T cells in the human immune response to cytomegalovirus. J. Clin. Invest. 103, 1437–1449 (1999).

19. Couzi, L., et al. Gamma-delta T cell expansion is closely associated with cytomegalovirus infection in all solid organ transplant recipients. Transpl. Int. 24, e40–e42 (2011).

20. Lafarge, X., et al. Cytomegalovirus Infection in Transplant Recipients Resolves When Circulating γδ T Lymphocytes Expand, Suggesting a Protective Antiviral Role. J. Infect. Dis. 184, 533–541 (2001).

21. Kaminski, H., et al. Characterization of a Unique γδ T-Cell Subset as a Specific Marker of Cytomegalovirus Infection Severity. J. Infect. Dis. 223, 655–666 (2021).

22. Chan, S., et al. Cytomegalovirus drives Vδ1+ γδ T cell expansion and clonality in common variable immunodeficiency. Nat. Commun. 15, 4286 (2024).

23. Willcox, C. R., et al. Cytomegalovirus and tumor stress surveillance by binding of a human γδ T cell antigen receptor to endothelial protein C receptor. Nat. Immunol. 13, 872–879 (2012).

24. Vermijlen, D., Gatti, D., Kouzeli, A., Rus, T. & Eberl, M. γδ T cell responses: How many ligands will it take till we know? Semin. Cell Dev. Biol. 84, 75–86 (2018).

25. Deseke, M., et al. A CMV-induced adaptive human Vδ1+ γδ T cell clone recognizes HLA-DR. J. Exp. Med. 219, e20212525 (2022).

26. Halary, F., et al. Shared reactivity of V{delta}2(neg) {gamma}{delta} T cells against cytomegalovirus-infected cells and tumor intestinal epithelial cells. J. Exp. Med. 201, 1567–1578 (2005).

27. Vermijlen, D., et al. Human cytomegalovirus elicits fetal γδ T cell responses in utero. J. Exp. Med. 207, 807–821 (2010).

28. Ravens, S., et al. Human γδ T cells are quickly reconstituted after stem-cell transplantation and show adaptive clonal expansion in response to viral infection. Nat. Immunol. 18, 393–401 (2017).

29. Wang, J., et al. Fetal and adult progenitors give rise to unique populations of CD8+ T cells. Blood 128, 3073–3082 (2016).

30. Tabilas, C., Smith, N. L. & Rudd, B. D. Shaping immunity for life: Layered development of CD8+ T cells. Immunol. Rev. 315, 108–125 (2023).

31. Ikuta, K., et al. A developmental switch in thymic lymphocyte maturation potential occurs at the level of hematopoietic stem cells. Cell 62, 863–874 (1990).

32. Vermijlen, D. & Prinz, I. Ontogeny of Innate T Lymphocytes – Some Innate Lymphocytes are More Innate than Others. Front. Immunol. 5, (2014).

33. Sanchez Sanchez, G., Tafesse, Y., Papadopoulou, M. & Vermijlen, D. Surfing on the waves of the human γδ T cell ontogenic sea. Immunol. Rev. 315, 89–107 (2023).

34. Tieppo, P., et al. The human fetal thymus generates invariant effector γδ T cells. J. Exp. Med. 217, (2020).

35. Papadopoulou, M., Sanchez Sanchez, G. & Vermijlen, D. Innate and adaptive γδ T cells: How, when, and why. Immunol. Rev. 298, 99–116 (2020).

36. Dimova, T., et al. Effector Vγ9Vδ2 T cells dominate the human fetal γδ T-cell repertoire. Proc. Natl. Acad. Sci. 112, E556–E565 (2015).

37. Papadopoulou, M., et al. Fetal public Vγ9Vδ2 T cells expand and gain potent cytotoxic functions early after birth. Proc. Natl. Acad. Sci. U. S. A. http://doi.org/10.1073/pnas.1922595117 (2020) doi:10.1073/pnas.1922595117.

38. Davey, M. S., et al. Human neutrophil clearance of bacterial pathogens triggers anti-microbial γδ T cell responses in early infection. PLoS Pathog. 7, e1002040 (2011).

39. Tan, L., et al. A fetal wave of human type 3 effector γδ cells with restricted TCR diversity persists into adulthood. Sci. Immunol. 6, eabf0125 (2021).

40. León-Lara, X., et al. γδ T cell profiling in a cohort of preterm infants reveals elevated frequencies of CD83+ γδ T cells in sepsis. J. Exp. Med. 221, e20231987 (2024).

41. Gattinoni, L., et al. A human memory T-cell subset with stem cell-like properties. Nat. Med. 17, 1290–1297 (2011).

42. Tsui, C. & Kallies, A. Unwrapping stemness to revive T cells. Science 386, 148–149 (2024).

43. Krishnan, S., et al. Amphiregulin-producing γδ T cells are vital for safeguarding oral barrier immune homeostasis. Proc. Natl. Acad. Sci. U. S. A. 115, 10738–10743 (2018).

44. du Halgouet, A., et al. Role of MR1-driven signals and amphiregulin on the recruitment and repair function of MAIT cells during skin wound healing. Immunity 56, 78–92.e6 (2023).

45. Kennedy-Crispin, M., et al. Human keratinocytes’ response to injury upregulates CCL20 and other genes linking innate and adaptive immunity. J. Invest. Dermatol. 132, 105–113 (2012).

46. Nielsen, H. V., et al. Nr4a1 and Nr4a3 redundantly control clonal deletion and contribute to an anergy-like transcriptome in auto-reactive thymocytes to impose tolerance in mice. Nat. Commun. 16, 784 (2025).

47. Liberzon, A., et al. Molecular signatures database (MSigDB) 3.0. Bioinformatics 27, 1739–1740 (2011).

48. Papadopoulou, M., et al. TCR Sequencing Reveals the Distinct Development of Fetal and Adult Human Vγ9Vδ2 T Cells. J. Immunol. 203, 1468–1479 (2019).

49. Sanchez Sanchez, G., et al. Identification of distinct functional thymic programming of fetal and pediatric human γδ thymocytes via single-cell analysis. Nat. Commun. 13, 5842 (2022).

50. Yang, Q., Safina, K. R. & Borcherding, N. scRepertoire 2: Enhanced and Efficient Toolkit for Single-Cell Immune Profiling. 2024.12.31.630854 Preprint at 10.1101/2024.12.31.630854 (2024).

51. Foss, A., Markatou, M., Ray, B. & Heching, A. A semiparametric method for clustering mixed data. Mach. Learn. 105, 419–458 (2016).

52. Jiang, L., et al. Overexpression of SMC4 activates TGFβ/Smad signaling and promotes aggressive phenotype in glioma cells. Oncogenesis 6, e301–e301 (2017).

53. Vander Ark, A., Cao, J. & Li, X. TGF-β receptors: In and beyond TGF-β signaling. Cell. Signal. 52, 112–120 (2018).

54. Li, M. O., Sanjabi, S. & Flavell, R. A. Transforming Growth Factor-β Controls Development, Homeostasis, and Tolerance of T Cells by Regulatory T Cell-Dependent and -Independent Mechanisms. Immunity 25, 455–471 (2006).

55. Oh, S. A. & Li, M. O. TGF-β: Guardian of T Cell Function. J. Immunol. 191, 3973–3979 (2013).

56. Rudd, B. D. Neonatal T Cells: A Reinterpretation. Annu. Rev. Immunol. 38, 229–247 (2020).

57. Vaaben, A. V., et al. In Utero Activation of Natural Killer Cells in Congenital Cytomegalovirus Infection. J. Infect. Dis. 226, 566–575 (2022).

58. Semmes, E. C., et al. Maternal Fc-mediated non-neutralizing antibody responses correlate with protection against congenital human cytomegalovirus infection. J. Clin. Invest. 132, (2022).

59. Semmes, E. C., et al. ADCC-activating antibodies correlate with decreased risk of congenital human cytomegalovirus transmission. JCI Insight 8, (2023).

60. Bodéus, M., et al. Human cytomegalovirus in utero transmission: follow-up of 524 maternal seroconversions. J. Clin. Virol. Off. Publ. Pan Am. Soc. Clin. Virol. 47, 201–202 (2010).

61. Enders, G., Daiminger, A., Bäder, U., Exler, S. & Enders, M. Intrauterine transmission and clinical outcome of 248 pregnancies with primary cytomegalovirus infection in relation to gestational age. J. Clin. Virol. Off. Publ. Pan Am. Soc. Clin. Virol. 52, 244–246 (2011).

62. Pass, R. F., Fowler, K. B., Boppana, S. B., Britt, W. J. & Stagno, S. Congenital cytomegalovirus infection following first trimester maternal infection: symptoms at birth and outcome. J. Clin. Virol. Off. Publ. Pan Am. Soc. Clin. Virol. 35, 216–220 (2006).

63. Revello, M. G., et al. Diagnosis and outcome of preconceptional and periconceptional primary human cytomegalovirus infections. J. Infect. Dis. 186, 553–557 (2002).

64. Beersma, M. F., Bijlmakers, M. J. & Ploegh, H. L. Human cytomegalovirus down-regulates HLA class I expression by reducing the stability of class I H chains. J. Immunol. Baltim. Md 1950 151, 4455–4464 (1993).

65. Sell, S., et al. Control of Murine Cytomegalovirus Infection by γδ T Cells. PLOS Pathog. 11, e1004481 (2015).

66. Kajubi, R., et al. Monthly sulfadoxine–pyrimethamine versus dihydroartemisinin–piperaquine for intermittent preventive treatment of malaria in pregnancy: a double-blind, randomised, controlled, superiority trial. The Lancet 393, 1428–1439 (2019).

67. Boeckh, M., et al. Optimization of Quantitative Detection of Cytomegalovirus DNA in Plasma by Real-Time PCR. J. Clin. Microbiol. 42, 1142–1148 (2004).

68. Henrich, T. J., Gallien, S., Li, J. Z., Pereyra, F. & Kuritzkes, D. R. Low-Level Detection and Quantitation of Cellular HIV-1 DNA and 2-LTR Circles Using Droplet Digital PCR. J. Virol. Methods 186, 68–72 (2012).

69. Auton, A., et al. A global reference for human genetic variation. Nature 526, 68–74 (2015).

70. Zhang, F., Kang, H. M. & Yan, Y. popscle: A suite of population scale analysis tools for single-cell genomics data. The Center for Statistical Genetics at the University of Michigan School of Public Health.

71. Neavin, D., et al. Demuxafy: improvement in droplet assignment by integrating multiple single-cell demultiplexing and doublet detection methods. Genome Biol. 25, 94 (2024).

72. McGinnis, C. S., Murrow, L. M. & Gartner, Z. J. DoubletFinder: Doublet Detection in Single-Cell RNA Sequencing Data Using Artificial Nearest Neighbors. Cell Syst. 8, 329–337.e4 (2019).

73. Babdor, J., et al. Coordinated variation in the immune systems and microbiomes of healthy humans is linked to tonic interferon states. 2024.11.27.625750 Preprint at 10.1101/2024.11.27.625750 (2024).

74. Zappia, L. & Oshlack, A. Clustering trees: a visualization for evaluating clusterings at multiple resolutions. GigaScience 7, giy083 (2018).

75. Subramanian, A., et al. Gene set enrichment analysis: A knowledge-based approach for interpreting genome-wide expression profiles. Proc. Natl. Acad. Sci. 102, 15545–15550 (2005).

76. Suo, C., et al. Dandelion uses the single-cell adaptive immune receptor repertoire to explore lymphocyte developmental origins. Nat. Biotechnol. 42, 40–51 (2024).

77. Finak, G., et al. MAST: a flexible statistical framework for assessing transcriptional changes and characterizing heterogeneity in single-cell RNA sequencing data. Genome Biol. 16, 278 (2015).

78. Wickham, H. Ggplot2: Elegant Graphics for Data Analysis. (Springer-Verlag New York, 2016).

79. Bunis, D. G., Andrews, J., Fragiadakis, G. K., Burt, T. D. & Sirota, M. dittoSeq: universal user-friendly single-cell and bulk RNA sequencing visualization toolkit. Bioinformatics 36, 5535–5536 (2021).

80. Wilke, C. ggridges: Ridgeline Plots in ‘ggplot2’. (2024).

81. Zeileis, A., et al. colorspace: A Toolbox for Manipulating and Assessing Colors and Palettes. J. Stat. Softw. 96, 1–49 (2020).

82. Wilkins, D. treemapify: Draw Treemaps in ‘ggplot2’. (2025).

83. Kassambara, A. ggpubr: ‘ggplot2’ Based Publication Ready Plots. (2023).

