## Supplementary Information for "Congenital CMV infection drives oligoclonal expansion of cytotoxic γδ T cells from early fetal progenitors"

| **Name** | **Sequence** | **Purpose** |
| --- | --- | --- |
| UL123 Forward | TCCCGCTTATCCTCRGGTACA | qRT-PCR for cCMV |
| UL 123 Reverse | TGAGCCTTTCGAGGASATGAA | qRT-PCR for cCMV |
| UL55 Forward | TGGGCGAGGACAACGAA | qRT-PCR and ddPCR for cCMV |
| UL55 Reverse | TGAGGCTGGGAAGCTGACAT | qRT-PCR and ddPCR for cCMV |
| UL55 Probe | TGGGCAACCACCGCACTGAGG | ddPCR for cCMV |
| 10X TCR Outer F | AATGATACGGCGACCACCGAGATCTACACTCTTTCC CTACACGACGCTC | First round of γδ TCR enrichment for scRNAseq |
| Cgamma outer R | CAAGAAGACAAAGGTATGTTCCAG | First round of γδ TCR enrichment for scRNAseq |
| Cdelta outer R | GTAGAATTCCTTCACCAGACAAG | First round of γδ TCR enrichment for scRNAseq |
| 10X TCR Inner F | AATGATACGGCGACCACCGAGATCT | Second round of γδ TCR enrichment for scRNAseq |
| Cgamma inner R | AATAGTGGGCTTGGGGGAAACATCTGCAT | Second round of γδ TCR enrichment for scRNAseq |
| Cdelta inner R | ACGGATGGTTTGGTATGAGGCTGACTTCT | Second round of γδ TCR enrichment for scRNAseq |

**Supplementary Table 1 – Primers and Probes.** All primers and probes and associated uses.

| **Marker-Fluorophore** | **Clone** | **Manufacturer** | **Concentration** |
| --- | --- | --- | --- |
| CD27 – BV421 | M-T271 | Biolegend | 1:25 |
| CD14 – APC-Cy7 | M5E2 | Biolegend | 1:50 |
| CD19 – APC-Cy7 | HIB19 | Biolegend | 1:50 |
| Live/Dead Near-IR | - | Thermo Fisher | 1:416 |
| CCR7 – PE | G043H7 | Biolegend | 3:50 |
| CD3 – BV 510 | UCHT1 | Biolegend | 1:50 |
| CD45RA – BV650 | HI100 | Biolegend | 1:25 |
| TCR Vδ2 – FITC | Immu389 | Beckman-Coulter | 1:10 |
| TCR Vγ9 – APC | B3 | Biolegend | 1:25 |
| CD16 – PE-Dazzle | 3G8 | Biolegend | 3:50 |
| TCR Vδ1 – PE-Cy7 | TS8.2 | Life Technologies | 3:50 |
| Perforin – BV421 | δG9 | BD | 3:50 |
| Granzyme B – AlexaFluor 700 | GB11 | BD | 1:25 |
| Ki67 – PE | B56 | BD | 1:5 |
| HLA-DR – BV711 | L243 | Biolegend | 1:25 |

**Supplementary Table 2 – Antibodies.** All antibodies used for flow cytometry.

**Supplementary Table 3 – Gene Sets.** Full list of genes in each named gene set used for module scoring.

| **Naïve/Immature** | *LRRN3, MAD1L1, BEX3, RHOH, RAC2, BCL11B, IKZF2, MRPS6, LRRC8D, CXCR3, RCSD1, CORO1A, SLC25A46, RGS18, LEF1, CHMP7, DCK, CHI3L2, RBL2, MCUB, NREP, TAF9B, ORAI2, SLC9A3R1, ACAP1, KDM5B, MMP25-AS1, NELL2, TCF7, SMC4, SLC25A5, GALNT2, CRLF3, ARHGDIB, TUBB, UBE2E2, DAAM1, ITGA4, MBP, CCDC69, AFF3, VIPR2, LTB, FCMR, SERINC5, PIK3IP1, SPINT2, CUX1, A1BG, PPP2R5C, SLC5A3, DBN1, PRDX2, FXYD2, SPIB, CD52, CCR7, AC119396.1, RALA, SNHG7, THEMIS, PIM2, EIF3E, IFT80, GSTO1, TOX, PRKCH, CDKN2D, MYO1G, TMSB10, SELL, MAL, AP1S2, STMN3, RIPOR2, RAB37, LINC02446, STK17A, TTC1, RCBTB2, MPST, CCNI, NSF, PECAM1, ACTN1, LAT, ITGA6, RASGRP2, C12orf57, RACK1, CD7, GRAMD1C, MGST2, CPNE7, GPR15, ZNF580, ITM2A, FYB1, AQP3, TESPA1, NDUFA12, LDLRAP1, FCER1G, ADSL, PITPNC1, MGLL, DCP2, CCDC50, GRINA, CD5, RTKN2, BIRC2, AIF1, GTF3A, EPB41, EEF2, TAGLN2, PSMA1, LCK, LRRC28, PSIP1, EVL, EEF1A1, MARCKSL1, NUCB2, HSPB1, GYPC, KLF2, TMIGD2, MME, C4orf48, CBX3, ELF1, FKBP1A, CHST2, RHOG, STMN1, RGS10, DGKA, TKT, SEPT9, MLLT11, CLDND1, ZNF683, PARP1, CXXC5, SIRPG, ODF2L, GSTK1, LDHB, TMEM123, AES, CDCA7, TMEM204, HIST1H2AC, ITM2C, FAM118A, CSTB, LAT2, GRSF1, BCL6, YWHAH, CAMK4, HMGA1, SEPT6, NOG, TCF12, CYSLTR1, H2AFZ, GNG4, ITGAL, PPP1R18, CYB5A, SOX4, MIR181A1HG, AHI1, SMIM24, SNAP47, RASSF1, LST1* |
| --- | --- |
| **Cytotoxic Lymphocyte – Vδ1** | *KLRC3, IFITM2, ITGB1, FGFBP2, PRF1, NKG7, CCL5, TGFBR3, GZMB, CTSW, FCGR3A, S1PR5, PLEK, EFHD2, PYHIN1, KLRF1, CST7, KLRG1, ADRB2, BTN3A2, CX3CR1, TBX21, APMAP, CLIC3, FGR, LINC00649, PRSS23, BIN2, C1orf21, GZMH, HLA-DPB1, CADM1, ABI3, AC116366.3, CARD16, MYO1F, C12orf75, STOM, CHST12, ADAM8, GNPTAB, CTSC, LITAF, PLEKHF1, PTGER2, RASGRP1, SYNE1, FLNA, KANSL1-AS1, ADGRG1, LPCAT1, LINC00944, RNF213, ZBP1, CMC1, TIGIT, HLA-DRB1, HLA-DPA1, CD74, HLA-DRB5, ZEB2, RARRES3, GZMM, ZBTB38, HLA-DRA, EOMES, NCR1, LYST, IL10RA, ITGB2, LGR6, SLA2, NFATC2, TTC38, PTPN4, IDH2, KIF21A, F2R, DOK2, HCST, AC243829.2, IRF1, CYTOR, SASH3, CD84, CD3G, IKZF3, RABGAP1L, BATF, LINC00402, ATP2B4, PSMB10, SLF1, SAMD3, CEP78, PRR5L, MFSD10, PTP4A2, PPP1R16B, ACTA2, ARHGAP25, IQGAP2, FCRL6, CD8A, KIAA1551, SH2D1B, PTGDR, CD63, FCRL3, CD8B, AOAH, LINC02384, LILRB1, SELPLG* |
| **Type 3 Immunity** | *JAML, S100A4, TC2N, IL7R, S100A6, KLRB1, NCR3, ALOX5AP, SLC4A10, CCR6, PRSS35, DPP4, CXCR6, CEBPD, GYG1, RORA, ME1, GZMK, BLK, ERN1, SESN1, SPOCK2, IL23R, FKBP11, GCHFR, IFNGR1, LCP1, CDC42EP3, CA2, LGALS3, EML4, SYTL2, LAPTM5, RORC, GPR65, CERK, IL4I1, PNP, GLRX, IL18RAP, PERP, CCDC107, OSTF1, PBXIP1, RGS2, PHACTR2, KIAA0319L, IFNG-AS1, ANXA2* |

| **Positive Regulation of T cell Proliferation** | *CD24, EBI3,LILRB2, TNFSF13B, HHLA2, CORO1A, TNFRSF13C, TMIGD2, IL23R, CLECL1P, CD55, DHPS, AGER, EFNB1, AIF1, EPO, ICOSLG, PTPN22, CCDC88B, PYCARD, CD274, ANXA1, NCKAP1L, CD209, HLA-A, HLA-DMB, HLA-DPA1, HLA-DPB1, HLA-E, HMGB1, HES1, IGF1, IGF2, IGFBP2, IL1A, IL1B, IL2, IL2RA, IL4, IL6, IL6ST, IL12B, IL12RB1, IL15, IL18, JAK2, LEP, LGALS9, MIR21, MIR30B, CD46, KITLG, NCK1, PNP, FOXP3, IL23A, SASH3, PPP3CA, PRKCQ, GPAM, ZP4, PTPRC, SELENOK, IL21, RPS3, CCL5, CCL19, XCL1, SHH, BMI1, RASAL3, SLAMF1, SLC7A1, SPN, SPTA1, STAT5B, SYK, TFRC, TGFBR2, TRAF6, TNFSF4, CCR2, TYK2, VCAM1, ZAP70, ZP3, VTCN1, PDCD1LG2, CD276, NCK2, CARD11, HAVCR2, TNFSF9, RIPK2, FADD, DNAJA3, CD1D, CD3E, CD6, CD28, CD80, CD86, CD40LG, CD70, CD81* |
| --- | --- |
| **T Cell Proliferation** | *CD24, TSPAN32, RASGRP1, EBI3, LILRB2, BTN2A2, GPNMB, CEBPB, TNFSF13B, LILRB1, MALT1, LILRB4, RIPK3, BTN3A1, GLMN, HHLA2, CORO1A, VSIG4, TNFRSF13C, CLC, TMIGD2, CR1, RC3H1, IL23R, CTLA4, CTNNB1, CTPS1, CLECL1P, CD55, DHPS, DLG1, AGER, DOCK2, EFNB1, AIF1, ELF4, EPO, ERBB2, FKBP1B, FOXJ1, TMEM131L, ICOSLG, NCSTN, SCRIB, CADM1, IL27, ABL1, FYN, IL4I1, LRRC32, PTPN22, PLA2G2D, TNFRSF21, CCDC88B, LGALS9B, PYCARD, CD274, ANXA1, NCKAP1L, PLA2G2E, CD209, HLA-A, HLA-DMB, HLA-DPA1, HLA-DPB1, HLA-DRB1, HLA-E, HLA-G, HMGB1, HES1, CLEC4G, IGF1, IGF2, IGFBP2, IHH, IL1A, IL1B, IL2, IL2RA, IL4, IL6, IL6ST, IL10, IL12B, IL12RB1, IL15, TNFRSF9, IL18, IDO1, IRF1, JAK2, ARG1, ARG2, LEP, LGALS3, LGALS9, LIPA, LMO1, MIR181C, MIR21, MIR30B, CD46, KITLG, MSN, NCK1, PNP, P2RX7, PAWR, FOXP3, ZBTB7B, IL23A, PIK3CG, PLA2G2A, PLA2G5, IL20RB, WNT4, SASH3, RC3H2, PPP3CA, PPP3CB, LMBR1L, PRKAR1A, PRKCQ, PRNP, CRTAM, PSMB10, TWSG1, PELI1, SH3RF1, GPAM, PTPN6, ZP4, PTPRC, BAX, SELENOK, RAC2, IL21, BCL6, RPS3, RPS6, CCL5, CCL19, BID, XCL1, SDC4, VSIR, SFTPD, PLA2G2F, SHH, BMI1, MARCHF7, RASAL3, SLAMF1, BMP4, SLC4A2, SLC7A1, LGALS9C, SLC11A1, SOS1, SOS2, SPN, SPTA1, STAT5B, SYK, PRDX2, TFRC, TGFBR2, TNFRSF1B, TP53, TRAF6, TNFSF4, CCR2, TNFRSF4, TYK2, SCGB1A1, VCAM1, ZAP70, ZP3, VTCN1, ARMC5, PDCD1LG2, CD276, NDFIP1, DOCK8, CASP3, ITCH, MAD1L1, NCK2, CARD11, HAVCR2, PDE5A, CBLB, TNFSF14, TNFSF9, TNFRSF14, RIPK2, FADD, CCND3, TNFSF18, SH2D2A, DNAJA3, CD1D, CD3E, CD6, DLG5, CD28, CD80, CD86, TNFSF8, MAPK8IP1, CD40LG, CD70, CD81, CD151* |

**Supplementary Table 4 – Clinical and demographic attributes.** Clinical and demographic attributes of individuals in this study.

|  | **cCMV+** | **cCMV-** |
| --- | --- | --- |
| Gestational age at delivery, weeks  *(Median and IQR)* | 39.9  *(39.1 - 40.4)* | 39.7  *(38.7- 40.8)* |
| Maternal age at delivery, years  *(Median and IQR)* | 22.6  *(18.8 – 24.6)* | 21.6  *(18.2 – 25.1)* |
| Sex | Female:  53.8% | Female:  50.7% |
| Placental malaria | Positive:  84.6% | Positive:  62.7% |
| Microcephaly, ≤3^rd^ percentile | 25% | 2.7% |
| Small for gestational age,  ≤3^rd^ percentile | 25% | 5.5% |

**Supplementary Table 5 – Expanded Clonotypes.** The top ten most expanded CDR3δ clonotypes by total frequency (# of cells) in all subjects. Paired gamma chain V gene call for the sequence is shown in the second column. Boxes are shaded if a neonate possessed that CDR3δ clonotype.

*The CALGELDDKLIF germline clonotype was previously described^27^.

† Contains zero N-additions.

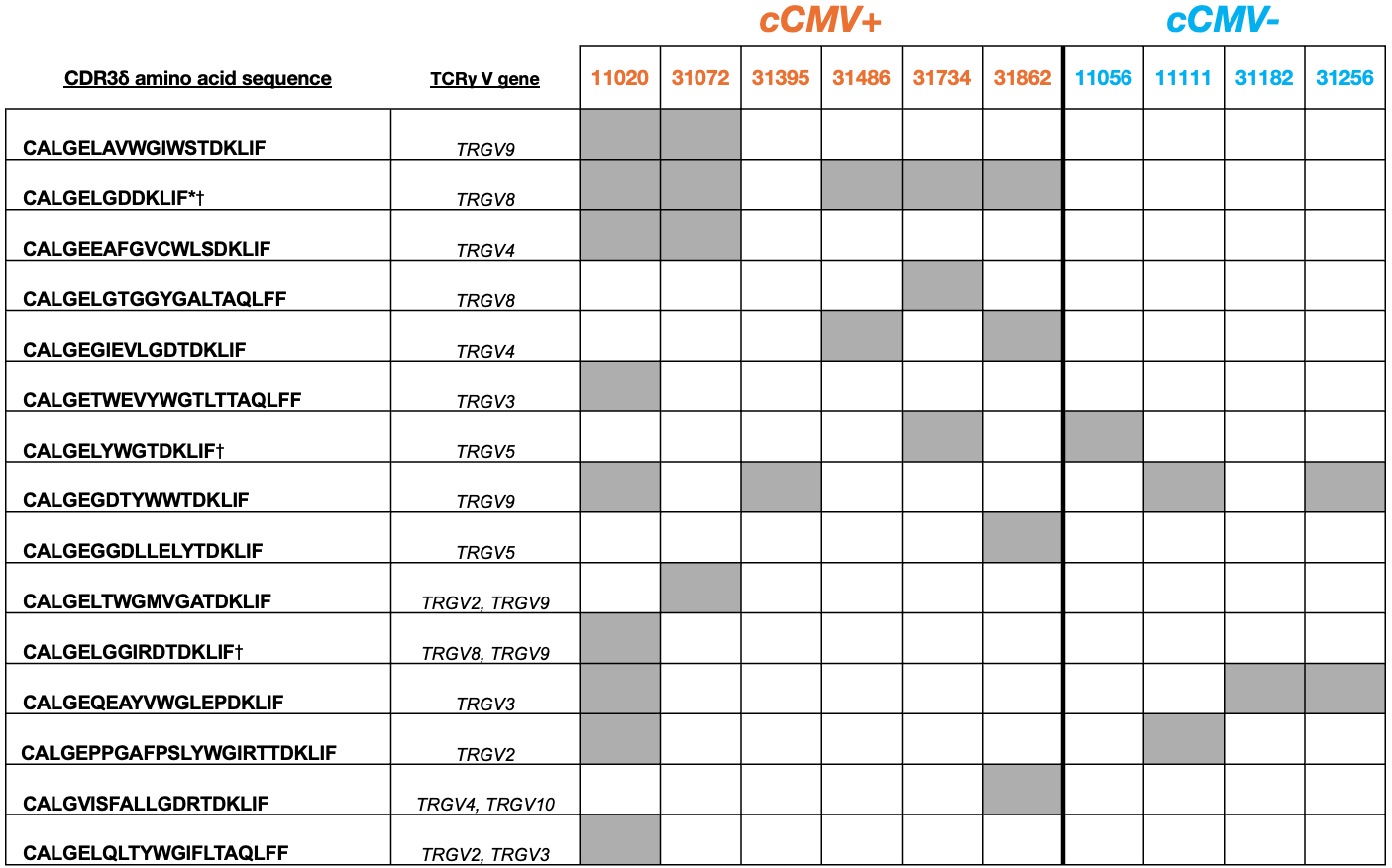

**Supplementary Figure 1. Flow cytometric gating strategies. A)** Gating strategy for γδ T cell subsets. **B)** Example flow gating of CD27^hi^ and CD27^lo/neg^ populations in total Vδ1+ γδ T cells in a representative cCMV- neonate (left) and cCMV+ neonate (right).

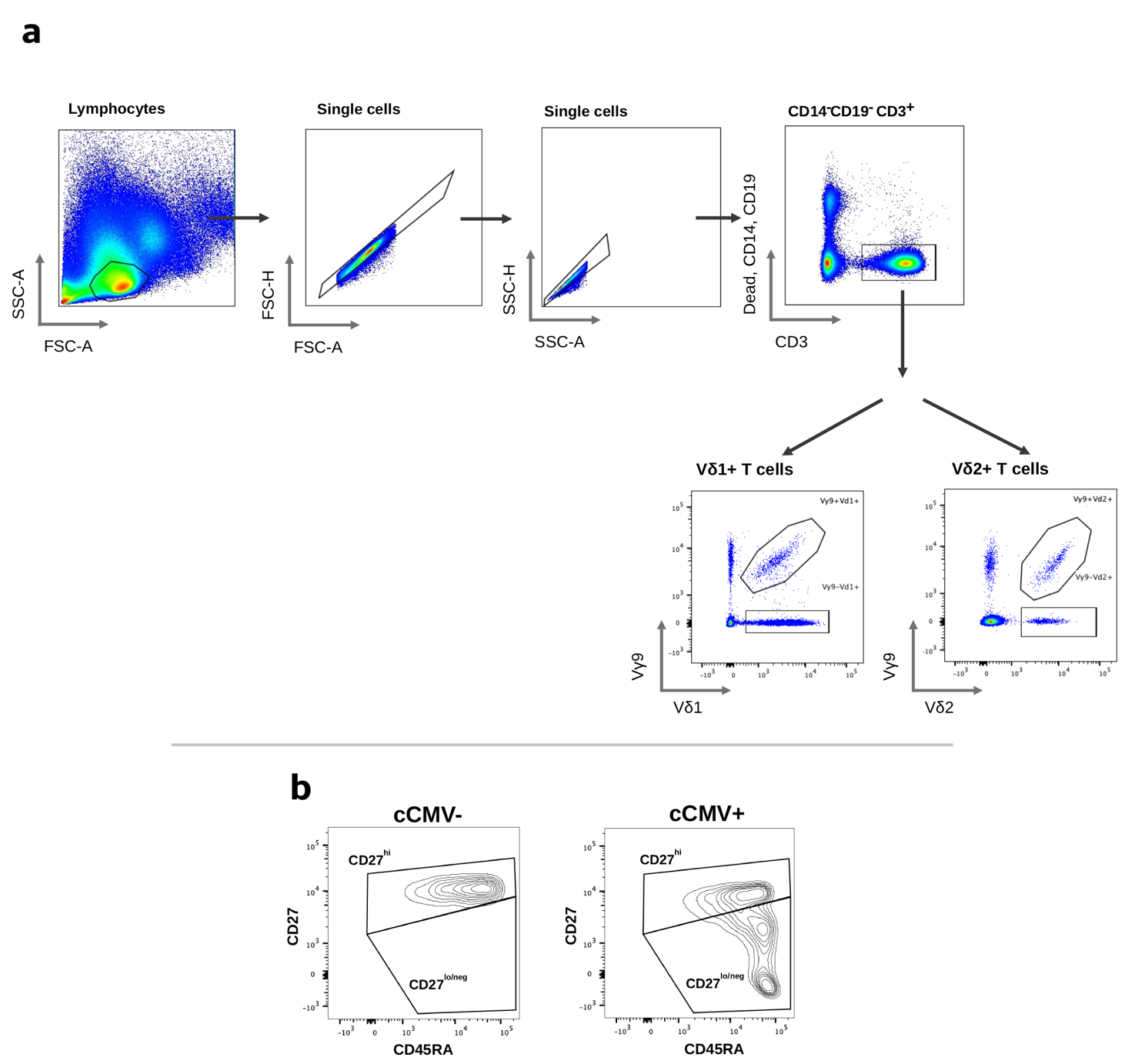

**Supplementary Figure 2. Clusters show differential enrichment in cCMV- and cCMV+ neonates. A)** Stacked barplots showing the proportion of cells in each cluster that come from cCMV- (blue) or cCMV+ (orange) infants. Asterisks denote a significant p-value ≤ 0.0001 for a Chi-square test comparing the observed distribution of cells for that cluster compared to the expected distribution of all cells. **B)** Boxplots displaying the frequency of cells that appear in each cluster per individual. Medians and IQR for cCMV- (blue) or cCMV+ (orange) neonates shown, each point is an individual. P-values were determined by Wilcoxon rank sum.

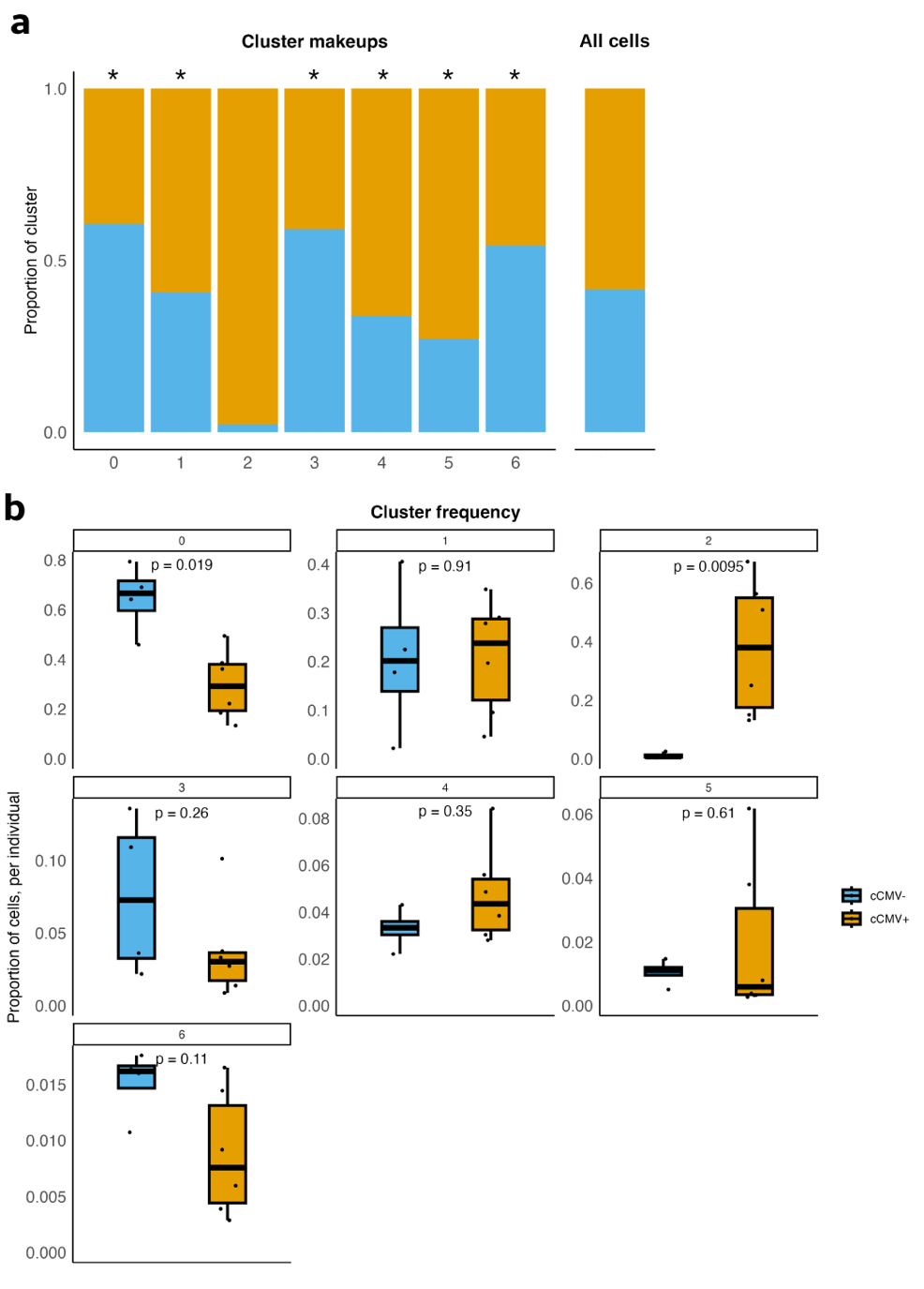

**Supplementary Figure 3. Tree maps of TCR repertoires from cCMV+ and cCMV- neonates. A)** Tree map representations of the TCRγ repertoires of cCMV+ (n = 6) or cCMV- (n = 4) individuals. **B)** Tree map representations of the TCRδ repertoires of cCMV+ (n = 6) or cCMV- (n = 4) individuals. For *A* and *B*, color scale indicates how many other individuals the TCR clonotype is shared with; each rectangle within a repertoire represents one clonotype and size of the rectangle corresponds to the proportion of the total repertoire that the clonotype occupies.

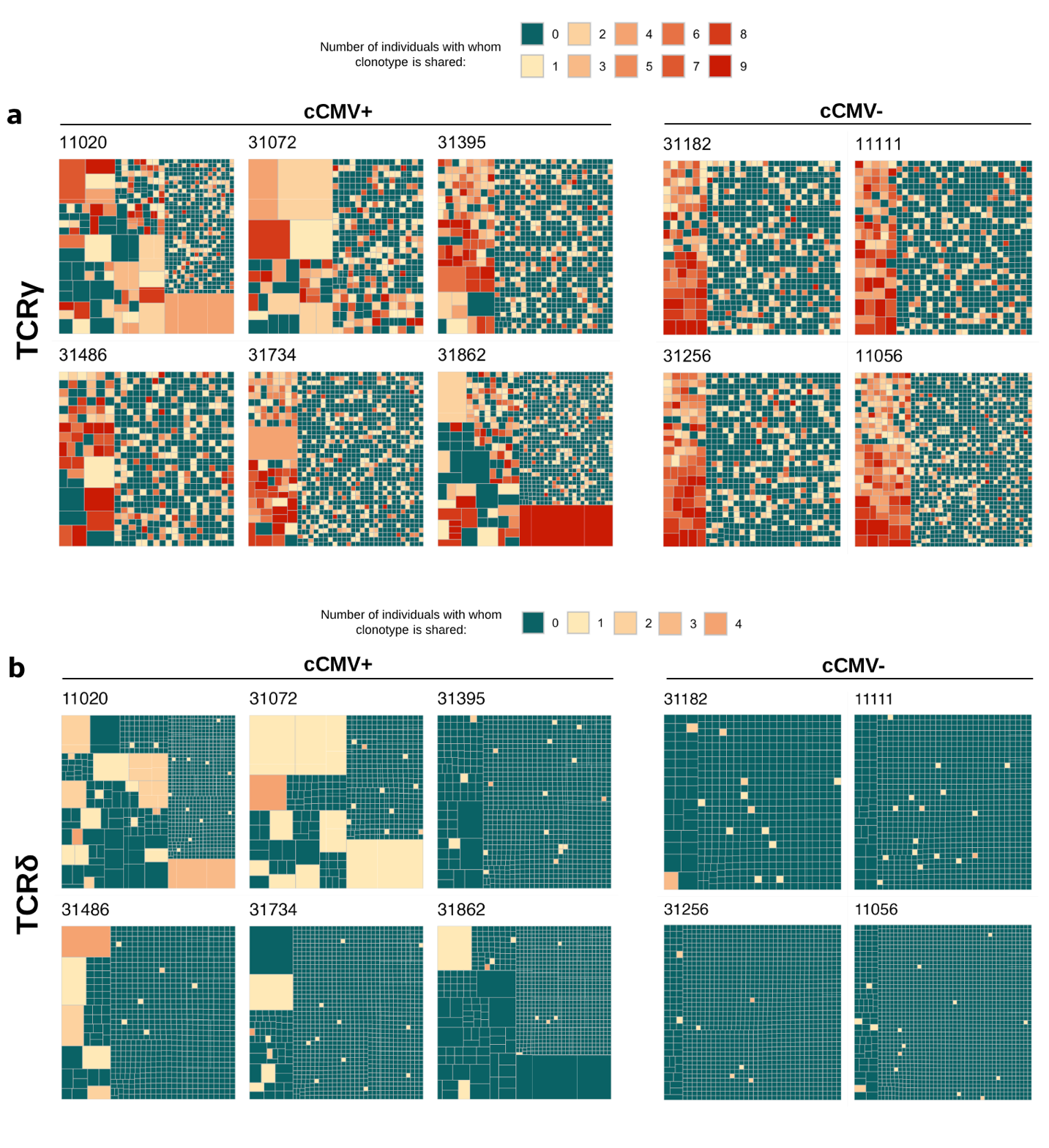
